# An integrated in silico-in vitro workflow for discovering high-affinity, selective antibodies to the KRAS(G12D)-MHC I complex

**DOI:** 10.1101/2025.08.31.673313

**Authors:** SangPhil Ahn, Tae-Sung Oh, Seonghyuk Suh, Joon-Young Jeon, Sangjoon Lah, Kyeongmin Ryu, Hyeonkyeong Kim, Eun Gyo Lee, Hyejin Lee, JooYeon Lee, Dong-Kyun Kim, Bo Mi Ku, Wooram Jung, Myung-Ju Ahn, Jae U. Jung, Yong-Sung Kim, Byung-Ha Oh, Bo-Seong Jeong

## Abstract

Antibodies that recognize peptide–loaded class I major histocompatibility complex (pMHC I) molecules could enable therapeutic targeting of intracellular oncogenic proteins, yet their discovery has been hampered by the small size of peptide antigens and allele-specificity. We describe an integrated in silico–in vitro workflow for generating high-affinity, selective antibodies to KRAS(G12D)_10_ presented by HLA-C*08:02, a clinically validated cancer neoantigen. In silico, multiple human antibody-derived variable fragments (Fvs) plausibly docked to the target pMHC were generated, followed by limited complementarity-determining region (CDR) sequence design. In vitro, CDR diversity was introduced at 3–4 positions per Fv to construct yeast surface display library for iterative selections. This workflow yielded antibodies with exclusive binding to KRAS(G12D)_10_/HLA-C*08:02 without cross-reactivity. Affinity maturation achieved nanomolar dissociation constants, and incorporation into chimeric antigen receptor T cells enabled specific activation against target-positive cells. This study establishes a practical design-to-function pipeline for TCR-like antibody discovery, and demonstrates the feasibility of therapeutic targeting against KRAS(G12D)-driven malignancies.

## Introduction

The Kirsten rat sarcoma viral oncogene homolog (KRAS) is one of the most frequently mutated proto-oncogenes in human cancers, with activating mutations at the glycine 12 (G12X) common in pancreatic, colorectal, and non–small cell lung cancers (1, 2). Although a range of small-molecule inhibitors, including sotorasib and adagrasib, have been developed for select alleles such as KRAS(G12C), these therapies are limited by allele specificity and often exhibit transient clinical benefits due to pathway reactivation or resistance (3–5). Additionally, broader strategies targeting pan-KRAS or other common variants, such as G12D and G12V, remain under intensive investigation but face challenges of specificity and therapeutic index (6, 7).

KRAS cannot be reached by conventional antibodies because it is intracellular, but peptide-loaded class I MHC (pMHC) displays fragments of intracellular proteins at the cell surface, providing a surface-exposed alternative target. (8). Leveraging this antigen-presentation axis, multiple therapeutic strategies have emerged, including cancer vaccines, T-cell receptor (TCR)–engineered therapies, and, more recently, pMHC-directed biologics such as TCR-like antibodies (9–11).

Among KRAS-derived neoantigens, the KRAS(G12D) decamer peptide (KRAS(G12D)_10_; GADGVGKSAL) presented by human leukocyte antigen C*08:02 (HLA-C*08:02) has emerged as a clinically validated target (12, 13). This epitope qualifies as a public antigen: a shared, tumor-specific peptide generated by recurrent driver mutations and displayed by a common HLA allele, and is therefore broadly actionable in defined patient subsets (14, 15). NT-112, a TCR-engineered T-cell therapy targeting KRAS(G12D)_10_/HLA-C*08:02, is in an ongoing phase I/II trial (NCT06218914), building on reports of durable responses in two patients with advanced pancreatic cancer treated under compassionate use (12, 13). These data position KRAS(G12D)_10_/HLA-C*08:02 among the most well-validated neoantigen-MHC complexes to date.

Despite this precedent, therapeutic targeting of pMHC complexes remains technically demanding. Soluble TCRs require laborious refolding and stabilization, and TCR-engineered T cells rely on ex vivo cell engineering, together imposing manufacturing burdens and limiting patient access (16, 17). In addition, cancer vaccines rely on the endogenous T-cell repertoire, which in advanced disease is often suboptimally primed or functionally exhausted (18). TCR-like antibodies, which combine TCR-level specificity with the manufacturability and stability of antibody scaffolds, represent an alternative modality amenable to broad deployment.

The development of antibodies that specifically recognize short peptides, typically 8-11 amino acids in length, presented on MHC remains a considerable challenge (19). Discrimination must be achieved on highly similar HLA scaffolds, where informative contacts are largely confined to solvent-exposed peptide side chains. Consequently, conventional screening often fails to recover such rare binders. Recent progress in computational design has yielded de novo binders against specified pMHC epitopes using diffusion-based generative models and structure-guided docking pipelines (20–23).

Here, we report an integrated in silico-in vitro approach to design and develop high-affinity TCR-like antibodies that selectively target the KRAS(G12D)_10_/HLA-C*08:02/β_2_-microglobulin (β2m) complex (hereafter called KRAS(G12D)/HLA-C8). This approach combines epitope-centric in silico design with focused complementarity-determining region (CDR) library screening to identify variants with strong binding affinity and exquisite specificity. We delineate the epitope-focused binding mode, confirm orthogonal specificity across related pMHCs, and demonstrate activity in a chimeric antigen receptor (CAR) T cell format. The framework is modular and supports broader development of pMHC-targeting antibodies, and underscores the translational relevance of the KRAS(G12D) neoantigen in HLA-C*08:02–positive tumors.

## Results

### Integrated in silico-in vitro approach to discover TCR-like antibodies to KRAS(G12D)/HLA-C8

Antibody-antigen interactions occur primarily through CDRS, which form flexible loops lacking defined secondary structures. This inherent structural plasticity poses challenges for computational antibody design, limiting generalizability and reducing success rates with current methods (24, 25). To address these limitations, we applied a previously developed pipeline that maintains the canonical backbone conformation of CDR loops while constraining residue design to structurally compatible positions, guided by position-specific consensus preferences (26).

We first aligned available TCR-like antibody/pMHC complexes to KRAS(G12D)/HLA-C8 (27). The original pMHC in each complex was then replaced with KRAS(G12D)/HLA-C8, resulting in various docking orientations of the TCR-like antibodies relative to the target. Subsequently, we superposed variable fragments (Fvs) from human antibodies onto the aligned TCR-like antibodies. Candidate Fvs were ranked by steric clashes, geometric distance metrics, and visual inspection to identify models with plausible docking geometry and peptide accessibility. (Fig. 1A).

**Fig. 1.**
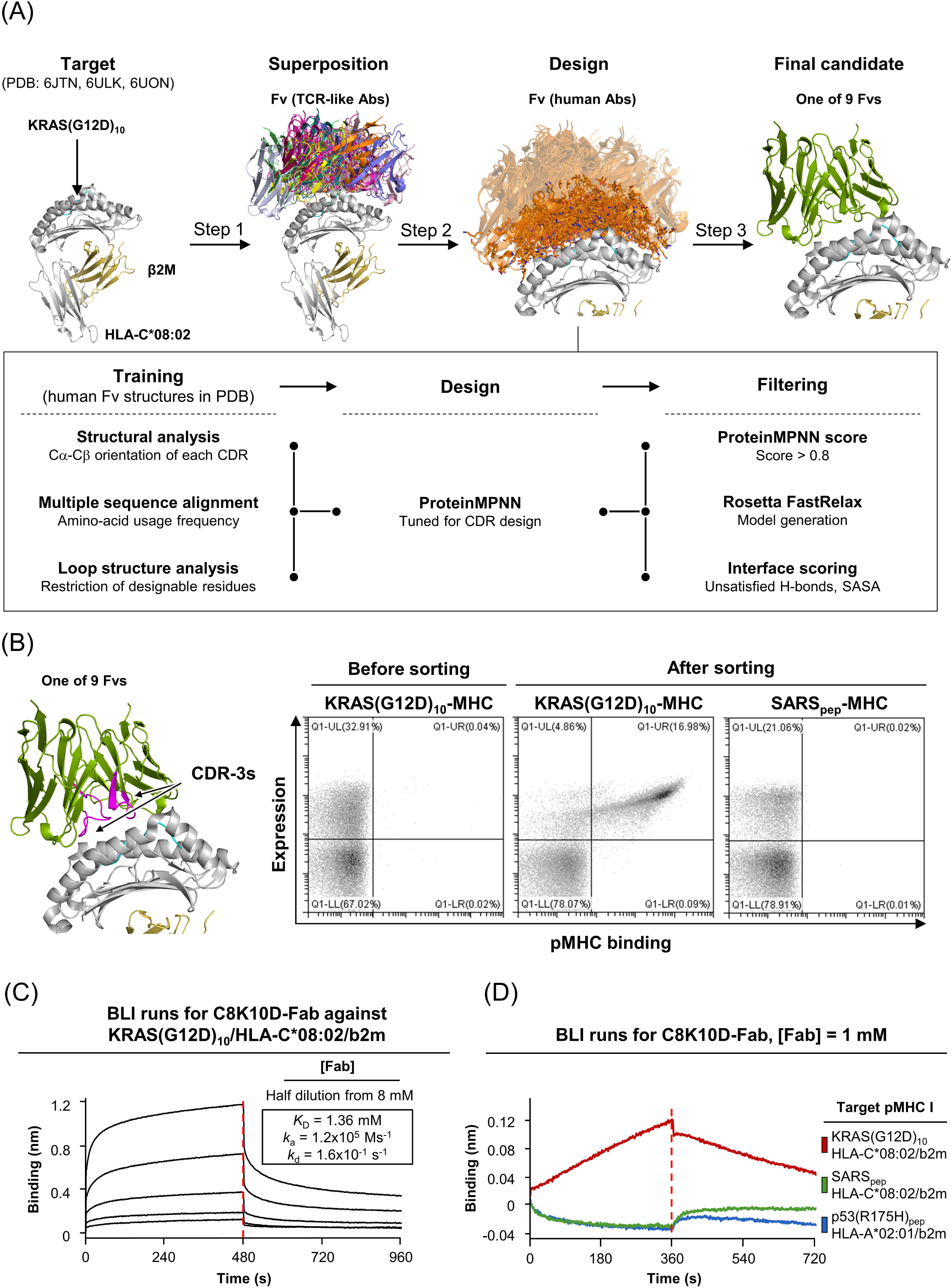
Computational antibody design integrated with experimental screening. **(A)** Computational antibody design was performed in three steps. (1) TCR-like antibody– pMHC complex structures were aligned to KRAS(G12D)_10_/HLA-C*08:02 (PDB: 6JTN, 6ULK, 6UON) to preserve docking geometry. (2) Human antibody Fv fragments were superposed onto the TCR-like frameworks. (3) Fvs were ranked by steric clash, geometric complementarity, and peptide accessibility. CDR-specific design was performed using a tuned ProteinMPNN model, with designable residues selected based on Cα–Cβ vector orientation and loop stability. Amino acid preferences were guided by MSA-based frequency. Designed sequences were filtered by ProteinMPNN score, visual inspection based on complex model, and interface features (SASA, H-bond satisfaction), yielding nine Fvs for library construction. **(B)** The designed variants underwent CDR-3s randomizing ≤ 4 residues to construct YSD library. The library was screened using fluorescently labeled KRAS(G12D)_10_/HLA-C*08:02 tetramers for positive selection and a tetramer of SARS_pep_/HLA-C*08:02 for negative selection by FACS. An enriched clone obtained through this process was designated C8K10D. **(C)** The binding affinity was quantified using BLI. The KRAS(G12D)_10_/HLA-C*08:02 complex was immobilized on streptavidin (SA) biosensor tips, followed by incubation with serial dilutions of C8K10D Fab starting at 8 μM and a dissociation phase. **(D)** The specificity of C8K10D was evaluated by BLI using KRAS(G12D)_10_/HLA-C*08:02 as the target pMHC, alongside SARS_pep_/HLA-C*08:02 and HLA-A*02:01 bound to HMTEVVRHC (human p53, residues 169-177) as control pMHCs. C8K10D Fab bound exclusively to the target pMHC.

To enable target-specific sequence optimization, we used an in-house version of proteinMPNN trained for antibody structures (28). Residue positions proximal to the peptide were identified based on structural analyses, including Cα–Cβ vector orientation and loop flexibility. Residues essential for preserving loop conformation were excluded from design to maintain native-like CDR backbone structures. Additionally, multiple sequence alignment (MSA)-based frequency profiles were applied to bias amino acid sampling toward evolutionarily preferred residues at each position (29, 30).

We initially filtered the design sequences based on ProteinMPNN scoring, then predicted structures using Rosetta FastRelax (31). Final candidates were evaluated for buried solvent-accessible surface area (SASA) and unsatisfied hydrogen bonds at the predicted binding interface. Based on these criteria, we selected nine Fvs for experimental evaluation.

To enhance diversity while preserving structural integrity, three to four residues within the CDR3 loops of each candidate were randomized, and the pooled inserts were cloned in scFv format into a yeast surface display vector and transformed into EBY100 to generate a single scFv yeast display library. This library was screened via magnetic-activated cell sorting (MACS) and iterative fluorescence-activated cell sorting (FACS). Positive selection used fluorescently labeled KRAS(G12D)/HLA-C8 tetramers. Specificity was monitored throughout the screening process by incorporating SARS_pep_ (ITDAVDCAL)/HLA-C*08:02 (SARS/HLA-C8) tetramers as a negative control during intermediate FACS rounds. Following multiple rounds of enrichment, a hit clone, C8K10D, was isolated (Fig. 1B and S1). We confirmed binding of the C8K10D Fab to the KRAS(G12D)/HLA-C8 by biolayer interferometry (BLI), with an equilibrium dissociation constant (*K*_D_) of ∼1.36 µM (Fig. 1C). No binding was detected against control pMHCs presenting unrelated peptides or alternative HLA alleles (Fig. 1D), demonstrating specificity and establishing C8K10D as a viable starting point for affinity maturation.

### Affinity enhancement via YSD

The initial lead binder, C8K10D, which exhibited a *K*_D_ of ∼1.36 µM, displayed binding affinity below the sub-100 nM threshold generally desirable for therapeutic antibody. To enhance binding affinity while maintaining target specificity, we constructed a secondary YSD library by randomizing the CDR-H1 and CDR-H2 regions. These regions were not diversified in the primary library but were structurally positioned to contribute to antigen engagement. To reduce avidity effects during selection, we performed the final FACS round with monomeric KRAS(G12D)/HLA-C8 complex (Fig. 2A and S2).

**Fig. 2.**
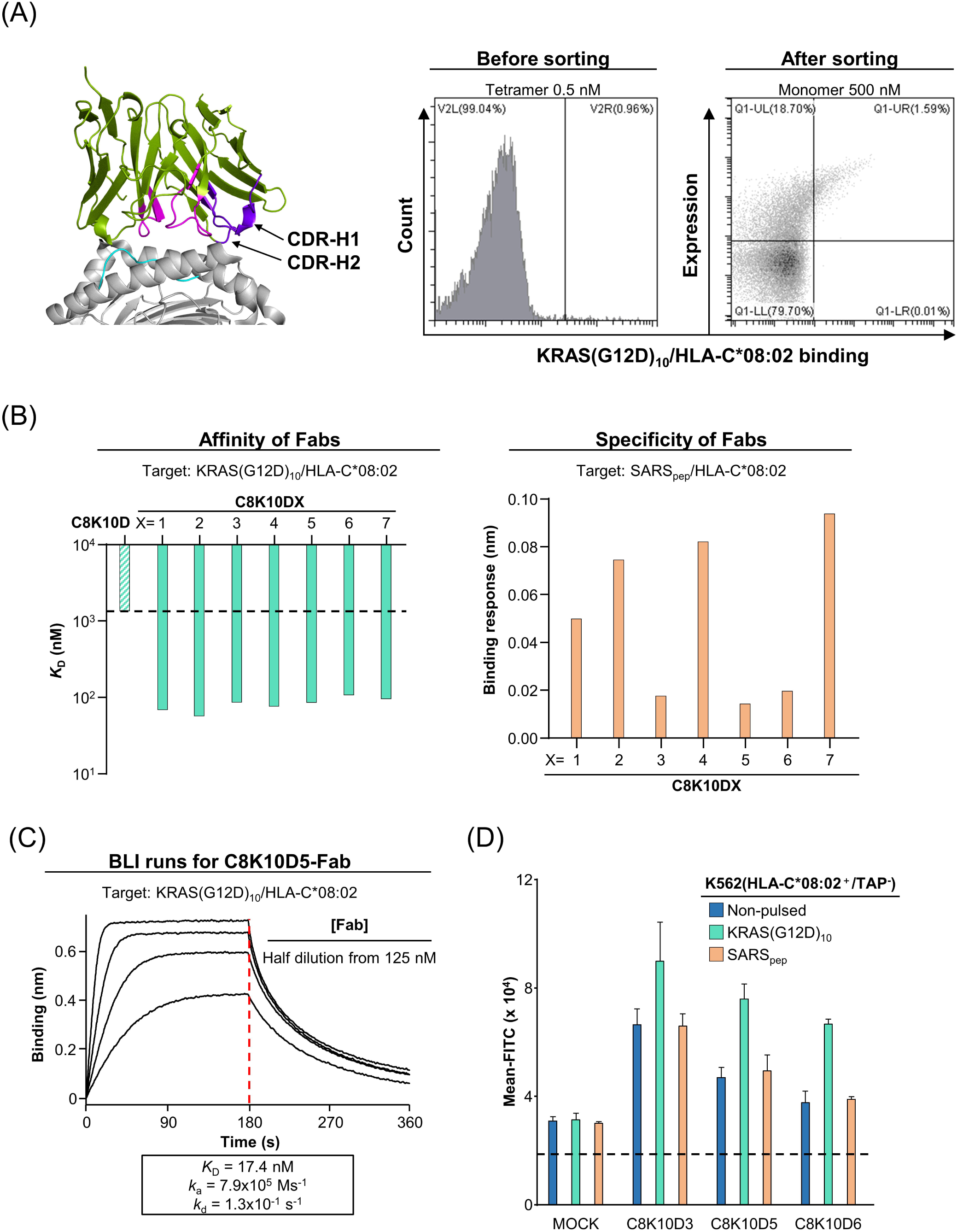
Binding affinity enhancement of C8K10D. **(A)** Secondary YSD library screening. The CDR-H1 and CDR-H2 regions of C8K10D were randomized to generate a secondary YSD library. MACS was used for initial enrichment, followed by isolation of seven clones (C8K10D1 - C8K10D7). **(B)** Assessment of target specificity and binding strength of the C8K10D variants. Binding of each Fab variant to the target pMHC was measured at a single concentration using BLI (left). Specificity was further evaluated using the control pMHC presenting SARS_pep_ peptide on HLA-C*08:02 (right). Among the variants, C8K10D5 and two additional clones exhibited no detectable binding to the control pMHC. **(C)** Measurement of binding affinity of C8K10D5 Fab. Serial dilutions of C8K10D5 Fab starting at 125 nM were incubated with KRAS(G12D)_10_/HLA-C*08:02 immobilized on SA sensor tips, followed by a dissociation phase. The *K_D_*, determined by fitting the sensorgrams, was 17.4 nM, which is markedly higher than that of the parental C8K10D Fab, which showed a *K_D_* of 1.36 µM (see Fig. 1C). **(D)** Cell-based binding assay. K562(HLA-C*08:02^+^/TAP^-^) cells pulsed with the KRAS(G12D)_10_ peptide or the SARS_pep_ peptide, or left unpulsed. Cells were then incubated with either C8K10D variants’ Fabs or control antibody (Mock), followed by flow cytometric analysis. All the variants showed increased binding signals in cells pulsed with the KRAS(G12D)_10_ peptide. However, the C8K10D5 Fab also displayed weak binding signals across all groups, indicating non-specific interaction with the cell surface.

Following iterative rounds of enrichment, seven distinct clones (C8K10D1 through C8K10D7) were isolated. Binding evaluation using BLI confirmed enhanced target binding across the panel, with several clones exhibiting nanomolar-range affinity to the KRAS(G12D)_10_/HLA-C*08:02 complex (Fig. 2B, left). In particular, three variants, C8K10D3, C8K10D5, and C8K10D6, demonstrated strong binding to the target pMHC while exhibiting minimal binding to a control pMHC presenting SARS_pep_ on the same HLA-C*08:02 allele (Fig. 2B, right).

We expressed these three clones as full-length immunoglobulin G (IgG) for further analysis. BLI measurements using serial dilutions confirmed that all three displayed dissociation constants in the range of approximately 20 nM, representing an approximately 70-fold improvement over the parental antibody, C8K10D (Fig. 2C and S3).

To assess cell-based binding, we employed K562 cells engineered to express HLA-C*08:02 and lack the transporter associated with antigen processing (TAP) expression, ensuring efficient surface presentation of exogenously loaded peptides. The cells were pulsed with either KRAS(G12D)_10_ peptide, SARS_pep_, or left unpulsed, then incubated with each antibody variant. Flow cytometric analysis revealed specific binding of all three antibodies to KRAS(G12D)_10_-pulsed cells, with markedly lower binding to SARS_pep_-pulsed or non-pulsed cells. However, low-level residual binding to control conditions was observed, exceeding background signals measured using a known TCR-like antibody specific for p53(R175H)/HLA-A*02:01 as a negative control (hereafter, control TCRL-Ab) (32) (Fig. 2D). These results prompted further engineering to eliminate off-target interactions.

### Mitigation of non-specific binding via cell-based selection

To address residual non-specific interactions with the cell surface, we investigated the feasibility of cell-based sorting using mammalian cells. Based on favorable expression yield and biophysical properties, we selected C8K10D5 as the lead candidate for this step. We assessed whether C8K10D5 could discriminate between target-presenting and non-target cells by co-culturing yeast-displayed C8K10D5 scFv with K562(HLA-C*08:02^+^/TAP^−^) cells either pulsed with KRAS(G12D)_10_ peptide or left unpulsed. Microscopic imaging revealed binding exclusively to peptide-pulsed K562 cells, confirming the mammalian-cell-based sorting can discriminate specific from non-specific binders (Fig. 3A and S4).

**Fig. 3.**
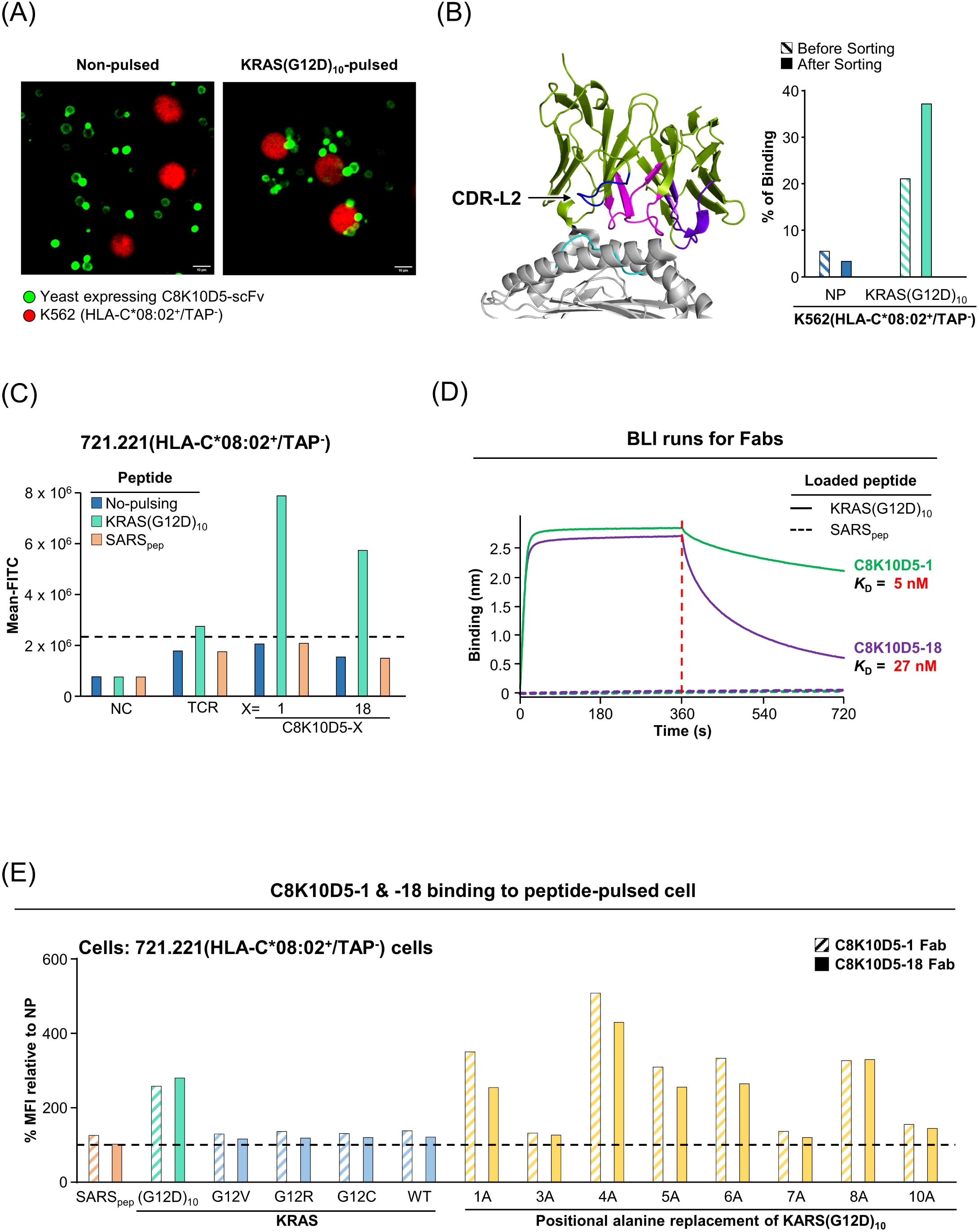
Elimination of non-specific binding of C8K10D5 and characterization of improved clones. **(A)** Representative confocal images of C8K10D5 scFv–displaying yeast co-incubated with K562(HLA-C*08:02^+^/TAP^-^) cells pulsed with KRAS(G12D)_10_ peptide or left unpulsed. Yeast cells were visualized with anti-Myc antibody staining (green), and target cells expressed cytoplasmic mScarlet-H (red). Specific binding of yeast was observed only with peptide-pulsed cells. Scale bar, 10 μm. **(B)** YSD library screening. Another yeast library was constructed in which the CDR-L2 region was randomized. Negative selection was performed using SARS_pep_-pulsed of non-pulsed mammalian cells (K562), and positive selection with fluorescently labeled KRAS(G12D)_10_/HLA-C*08:02 tetramer or monomer (Supplementary Fig 5). Incubation of enriched yeast clones with K562 cells revealed a reduction in non-specific binding from 5.51 % to 3.37 %. Binding to fluorescently labeled KRAS(G12D)_10_/HLA-C*08:02 increased from 21.13 % to 37.17 %, indicating both enhanced specificity and affinity. Two enriched clones were selected and designated C8K10D5-1 and C8K10D5-18. **(C)** Specificity of C8K10D5-1 and C8K10D5-18 was assessed by flow cytometry using peptide-pulsed 721.221(HLA-C*08:02^+^/TAP^-^) cells. Both Fab clones bound only to cells pulsed with the KRAS(G12D)_10_ peptide, while no binding was observed in cells pulsed with a control peptide or left unpulsed. Compared to a known TCR-like antibody specific for the target pMHC, the Fab clones demonstrated higher binding signals. **(D)** Binding affinity and specificity of C8K10D5-1 and C8K10D5-18 were evaluated by BLI. Single-concentration kinetics curves are shown. Kinetic analyses using multiple analyte concentrations yielded *K*_D_ values of 5 nM for C8K10D5-1 and 27 nM for C8K10D5-18. Neither clone exhibited detectable binding to control pMHCs. **(E)** Peptide specificity. Peptides incorporating alanine-scanning substitutions at each position of the 10-mer KRAS(G12D) peptide were used. 721.221(HLA-C*08:02^+^/TAP^-^) cells were pulsed with these alanine-substituted peptides and analyzed by flow cytometry. Both C8K10D5-1 Fab and C8K10D5-18 Fab exhibited loss of binding to specific alanine-substituted variants, confirming residue-level specificity. No binding was observed to the corresponding wild-type peptide, a control peptide, or other common KRAS G12 mutations, including G12V, G12C and G12R.

Following these observations, we constructed a sub-library of C8K10D5 by randomizing the CDR-L2 loop, which was predicted to contribute less to antigen recognition (Fig. 3B, left). Library screening was performed using negative selection against non-pulsed K562 cells and positive selection with KRAS(G12D)/HLA-C8 in the presence of non-target pulsed or non-pulsed K562 cells (Fig. S5). After sorting, yeast binding to non-pulsed K562 cells decreased from 5.51% to 3.37%, whereas binding to KRAS(G12D)_10_-pulsed cells increased from 21.13% to 37.17% (Fig. 3B, right and S6). Two enriched clones were isolated and designated as C8K10D5-1 and C8K10D5-18.

To evaluate specificity at the cellular level, we performed Fab binding assays on 721.221(HLA-C*08:02^+^/TAP^−^) cells pulsed with either KRAS(G12D)_10_, SARS_pep_, or no peptide as a control. While the K562 cells were previously used for sorting C8K10D Fab clones and initial functional assays, we utilized 721.221 cell line to confirm whether the observed specificity extends beyond a single cell line. 721.221(HLA-C*08:02^+^/TAP^−^) cells were generated following the same protocol applied to K562(HLA-C*08:02^+^/TAP^−^) cells, including peptide pulsing, ensuring consistent HLA expression, TAP deficiency, and peptide loading. As a positive control, we included a recombinant TCR previously reported to bind selectively to KRAS(G12D)/HLA-C8 (27). Both C8K10D5-1 and -18 Fabs bound selectively to KRAS(G12D)_10_-pulsed cells, showing no detectable binding to non-pulsed or SARS_pep_-pulsed cells (Fig. 3C). BLI-based kinetic analysis further revealed dissociation constants of 5 nM and 27 nM for C8K10D5-1 and C8K10D5-18, respectively, with no measurable binding to SARS_pep_-MHC (Fig. 3D).

We next assessed fine epitope specificity using 721.221(HLA-C*08:02^+^/TAP^−^) cells pulsed with peptides harboring alanine substitutions at each residue position of KRAS(G12D)_10_, and other mutants of G12 position in KRAS. Flow cytometric analysis demonstrated strong binding of both Fabs to the pulsed cells presenting KRAS(G12D)_10_, but no detectable interaction with alanine-substituted peptides at positions 3, 7, or 10, indicating that these positions are critical contact residues. No cross-reactivity was observed with the wild-type KRAS peptide, SARS_pep_, or other G12 mutant peptides (Fig. 3E and S7), confirming the residue-level specificity of the engineered variants.

### Structure-guided engineering for improved productivity and affinity

To further improve the biophysical properties of the lead candidate, we determined the co-crystal structure of C8K10D5-1 Fab in complex with the KRAS(G12D)/HLA-C8 using X-ray crystallography (Fig. 4A and Table 1). Although the resolution was limited to 4.5 Å, the overall binding orientation was clearly resolved and closely matched the design model (Figure 1A), adopting a diagonal docking geometry characteristic of TCR-like antibodies. Structural analysis identified surface-exposed residues in C8K10D5-1 that were not directly involved in antigen binding but we hypothesized that they contribute to contribute to protein stability.

**Fig. 4.**
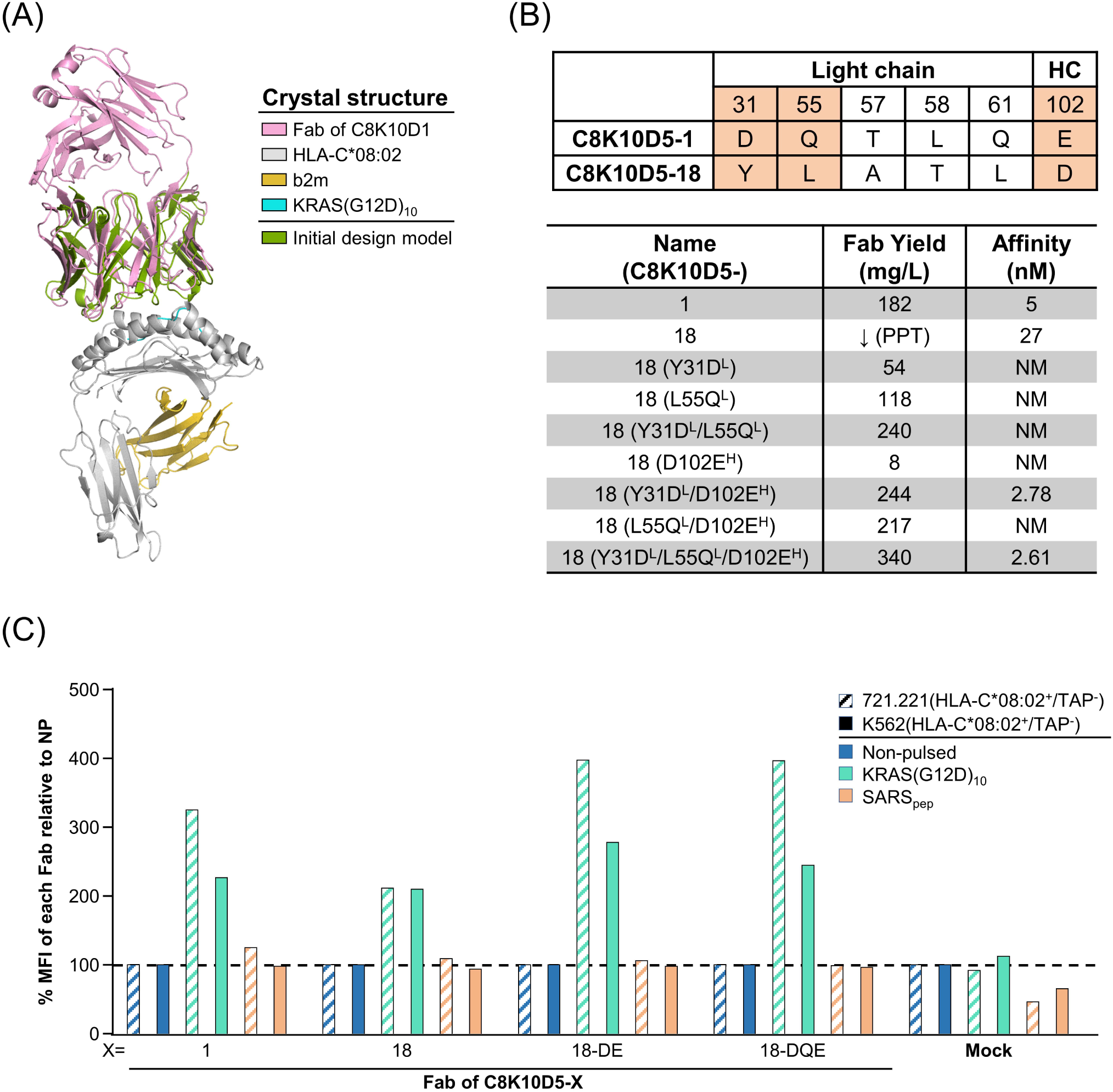
Structural characterization and structure-guided optimization of C8K10D5-18. **(A)** Crystal structure of C8K10D5-1 Fab bound to KRAS(G12D)_10_/HLA-C*08:02. The overall binding orientation closely matched the original design model, with a minor lateral displacement. The Fab docks onto the pMHC in a canonical diagonal orientation, forming a continuous interface across both the peptide and the α1/α2 helices of the HLA molecule. **(B)** Structure-guided engineering. To improve the solubility of C8K10D5-18, which is prone to precipitation, the indicated residues were substituted based on the crystal structure of C8K10D5-1 Fab/KRAS(G12D)_10_/HLA-C*08:02. The resulting variants exhibited enhanced solubility, with the best variant remaining soluble at 340 mg/L after purification. The C8K10D5-18 variants with improved production yield were evaluated for binding kinetics using BLI at a single concentration, confirming no detectable interaction with control pMHC I. Among the tested mutations, C8K10D5-18-DE and C8K10D5-18-DQE exhibited markedly reduced dissociation rates. Subsequent affinity measurements using serial dilutions and global kinetic fitting yielded *K*_D_ values of approximately 2.7[nM for both variants. **(C)** Specificity profiling using peptide-pulsed antigen-presenting cell lines. Fab binding was evaluated on K562(HLA-C*08:02^+^/TAP^-^) cells and 721.221(HLA-C*08:02^+^/TAP^-^) cells pulsed with either KRAS(G12D)_10_, a control peptide (SARS_pep_), or left unpulsed. C8K10D5-1, C8K10D5-18, and its engineered variants (C8K10D5-18DE and C8K10D5-18DQE) bound selectively to cells pulsed with the KRAS(G12D)_10_ peptide. A previously characterized TCR-like antibody served as a negative control (Mock).

**Table 1.** Data collection and refinement statistics.

| **Data Collection** | PDB ID: 9WJ6 |
| --- | --- |
| Space group | *P6<sub>5</sub>22* |
| Unit cell dimensions |  |
| a, b, c (Å) | 164.98, 164.98, 426.14 |
| $\alpha$, $\beta$, $\gamma$ (°) | 90.0, 90.0, 120.0 |
| Wavelength (Å) | 1.0000 |
| Resolution (Å) | 29.79-4.54 (4.702-4.54)<sup>a</sup> |
| R-merge | 0.4129 (5.189)<sup>a</sup> |
| I/$\sigma$(I) | 16.24 (5.51)<sup>a</sup> |
| Completeness | 98.32 (91.91)<sup>a</sup> |
| Redundancy | 38.5 (37.0)<sup>a</sup> |
| **Refinement** |  |
| No. of reflections | 20,329 (1,830)<sup>a</sup> |
| R<sub>work</sub>/R<sub>free</sub> (%) | 27.2/29.6 |
| R.m.s deviations |  |
| Bond (Å)/Angle (°) | 0.003/0.67 |
| Average B-values (Å<sup>2</sup>) | 169.01 |
| Ramachandran plot (%) |  |
| Favoured/Additionally allowed | 97.29/2.71 |
| Outliers | 0.0 |

<sup>a</sup>The numbers in parentheses are the statistics from the highest resolution shell.

Based on this analysis, we introduced three mutations, Y31D and L55Q in the light chain and D102E in the heavy chain, into the C8K10D5-18 backbone to generate engineered variants C8K10D5-18-DE (Y31D^L^/D102E^H^) and C8K10D5-18-DQE (Y31D^L^/L55Q^L^/D102E^H^). These substitutions led to marked improvements in Fab solubility, increasing the purified yield from undetectable levels to 217 mg/L and 340 mg/L, respectively. Moreover, BLI revealed enhanced binding affinities of 2.78 nM for C8K10D5-18-DE and 2.61 nM for -DQE (Fig. 4B and S8). Although these residues do not directly contact the pMHC, they likely contribute to favorable structural stabilization of the antibody framework.

To assess target specificity, we performed flow cytometric analysis using both K562(HLA-C*08:02^+^/TAP^-^) and 721.221(HLA-C*08:02^+^/TAP^-^) cells pulsed with KRAS(G12D)_10_ or SARS_pep_ peptides. Staining with Fabs of C8K10D5-1, -18, -18-DE, and - 18-DQE revealed that the engineered variants exhibited markedly increased binding to KRAS(G12D)_10_-pulsed cells relative to their parental clones, with minimal background staining of non-pulsed or SARS_pep_-pulsed cells. A previously described control TCRL-Ab showed no detectable binding (32) (Fig. 4C and S9). These results confirm that structure-guided optimization enhanced both the manufacturability and functional performance of C8K10D5-18 derivatives.

### Functional validation as chimeric antigen receptor (CAR)

To assess antigen-specific activation, we generated second-generation CAR constructs incorporating each scFvs of C8K10D5-1, -18, -18-DE, and -18-DQE fused to a CAR signaling cassette (Fig. 5, Top). Primary human T cells were engineered with the CAR constructs of the four distinct scFvs to generate CAR-T cells (Fig. S10A). These CAR-T cells were co-cultured with K562(HLA-C*08:02^+^/TAP^-^) or 721.221 (HLA-C*08:02^+^/TAP^-^) target cells pulsed with KRAS(G12D)_10_ peptide, with SARS_pep_-pulsed and unpulsed targets serving as controls to evaluate cytolytic potential.

**Fig. 5.**
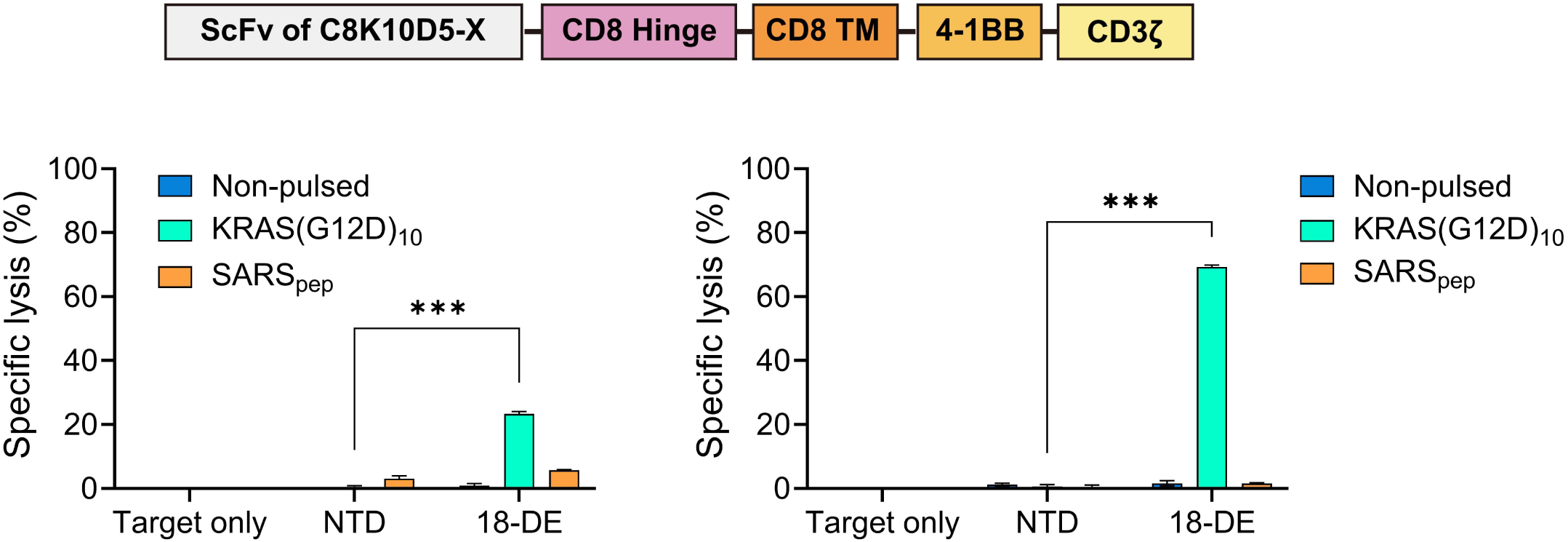
Antigen-specific activation of T cells by C8K10D5 variant-derived CARs. Top: Schematic of the chimeric antigen receptor (CAR) construct used to engineer Jurkat T cell (Top). The scFv derived from each C8K10D5 variant was fused to CD8α hinge and transmembrane domains, followed by 4-1BB and CD3ζ signaling domains. “GS” indicates a Gly-Ser linker, “TM” denotes the transmembrane region. Bottom: Peptide-dependent cytotoxicity of C8K10D5-18-DE–derived CAR-T cells. K562(HLA-C*08:02^+^/TAP^-^) cells and 721.221(HLA-C*08:02^+^/TAP^-^) target cells were pulsed with KRAS(G12D)_10_ or SARS_pep_ peptides or left unpulsed, and co-cultured with T cells expressing CAR derived from C8K10D5-18-DE at an effector-to-target ratio of 1:1 for 4 h. Cytotoxicity was measured by calcein-AM release assay. NTD, non-transduced T-cell control. Data are represented mean ± SD of triplicates; statistical analyses were performed using GraphPad Prism.

In a calcein-AM release assay, all CAR-T variants showed robust, peptide-specific lysis of KRAS(G12D)_10_-pulsed targets, whereas SARS_pep_-pulsed, target-only controls and non-transduced T cells exhibited minimal lysis (Fig. S10B). Among the variants, C8K10D5-18-DE exhibited the highest cytotoxicity (Fig. 5, Bottom). Taken together, these findings demonstrate that C8K10D5 variant–derived CARs elicit peptide-specific activation and cytotoxicity, preserving TCR-like recognition of pMHC complexes.

## Discussion

T cell receptor (TCR)-based therapeutics have emerged as a powerful class of cancer immunotherapies, with several modalities demonstrating clinical success. Tebentafusp, a bispecific T cell engager that combines a recombinant TCR with an anti-CD3 single-chain variable fragment, has been approved by the U.S. Food and Drug Administration (FDA) for the treatment of melanoma (33). More recently, the TCR-engineered T cell therapy afami-cel has also received FDA approval for synovial sarcoma (34). In parallel, de novo-designed proteins targeting pMHC complexes have emerged, enabled by advances in artificial intelligence (AI)-driven protein design (20–23). These include highly stable α-helical bundle binders generated by integrating generative backbone design with deep learning–based sequence optimization. However, these binders are structurally distinct from conventional biologics and may require new manufacturing or formulation strategies. In contrast, the antibody-based format described here represents a familiar scaffold for clinical development, compatible with established manufacturing platforms and regulatory pathways.

The TCR-like antibody identified in this study exhibits markedly higher affinity than typical TCRs. While TCRs generally have an *K*_D_ in the micromolar range, our lead candidate demonstrated the *K*_D_ value in the low nanomolar range and maintained strict specificity for the KRAS(G12D)_10_ peptide presented by HLA-C*08:02, without detectable cross-reactivity to closely related peptides. The strong target discrimination is especially notable because the peptide epitope differs from the wild-type sequence by only a single amino acid. Functional testing in CAR–modified T cells confirmed antigen-dependent activation, suggesting that this antibody could serve as the basis for off-the-shelf cell-based therapies. Given that tumor-specific antigens such as KRAS(G12D) are shared across multiple patients, such reagents have potential to enable broadly applicable T cell–engaging biologics.

Additionally, the CAR-T results suggest that this approach could be applied in alternative TCR-like cell therapy formats such as Synthetic TCR and Antigen Receptor T cells (STAR-T) and T cell Receptor Fusion Construct T cells (TruC) (35, 36). These formats, which incorporate antibody-based recognition into chimeric receptors that mimic endogenous TCR signaling, could be platforms for our candidates to exploit sensitivity of TCR pathways for detecting low-abundance targets. Beyond adoptive cell therapies, such candidate antibodies could be engineered into bispecific formats to redirect endogenous T cells toward tumor cells presenting the neoantigen, expanding therapeutic accessibility without the need for patient-specific manufacturing.

Developing antibodies against pMHC targets remains technically challenging because the antigenic surface is dominated by the MHC helices, while the relevant discriminating features reside within the bound peptide (37). For neoantigens, these features may involve a single mutated residue among a short 9-10 amino acid sequence, making the epitope one of the most demanding in terms of specificity (14). Computational antibody design is complicated further by the conformational plasticity of the CDR loops, which often limits prediction accuracy (24). Previous studies have reported limited success in loop-targeted design, but our approach minimized conformational uncertainty by restricting sequence design to structurally compatible positions while preserving canonical loop backbones (26). Incorporating experimental screening early in the process enabled identification of binders with the desired specificity and affinity. The crystal structure of the lead antibody–antigen complex validated the accuracy of the design model and revealed key features underlying its high specificity.

To enhance the screening process, we implemented a cell-based sorting method in which yeast-displayed antibodies were co-cultured with mammalian target cells. Previous studies have described mammalian cell co-culture–based sorting strategies (38, 39), but here we refined this approach by exercising stringent control over exogenous experimental conditions. By performing negative selection against non-pulsed cells and positive selection with peptide-pulsed cells under identical conditions, we efficiently eliminated binders with non-specific cell-surface interaction. This approach could be adapted to screen against other challenging targets, including membrane proteins that are difficult to purify in recombinant form. Moreover, by incorporating appropriate reporter systems into the mammalian target cells, this could enable sorting for binders that induce downstream signaling, thereby enriching for functionally active antibodies at an early stage.

Collectively, these results establish a generalizable design-to-function pipeline for generating high-affinity, high-specificity pMHC-targeting antibodies. When applied to the clinically validated KRAS(G12D) neoantigen, this approach yielded a promising therapeutic lead for HLA-restricted cancer immunotherapy. This framework can accelerate the development of precision biologics for a broad range of intracellular oncogenic targets presented by MHC molecules.

## Materials and Methods

### Computational design of TCR-like antibody

Antibody design was performed through a multi-step structural and sequence-based in silico workflow targeting the KRAS(G12D)_10_/HLA-C*08:02 complex (PDB: 6JTN, 6ULK, 6UON). To preserve plausible docking geometry, ten TCR-like antibody–pMHC I complex structures (PDB IDs: 1W72, 3CVH, 3GJF, 3HAE, 4WUU, 6UJ9, 6W51, 7BBG, 7RE7, and 7TR4) were structurally aligned to 6ULK using PyMOL (The PyMOL Molecular Graphics System, Version 4.6.0 Schrödinger, LLC.). Fv domains from human antibodies were then superposed onto each framework using backbone alignment, and candidates were ranked based on steric clash counts, peptide accessibility, and geometry consistency.

Sequence design was conducted using a custom-trained version of ProteinMPNN optimized for antibody CDRs (28). Designable residues were pre-selected based on structural criteria, including side-chain vector orientation (Cα–Cβ) toward the peptide and loop stability inferred from backbone torsion angles. Positions critical for maintaining canonical CDR loop conformations were excluded. Amino acid frequencies at each position were weighted using position-specific frequency profiles derived from multiple sequence alignments of human antibody repertoires.

Initial design filtering was performed using the ProteinMPNN score, retaining only sequences with scores >0.8 (28). Remaining sequences were modeled using Rosetta FastRelax (31). Final models were evaluated for buried SASA and unsatisfied hydrogen bonds at the antigen–antibody interface. A single design was selected for library construction based on the highest composite ranking across all metrics.

### Yeast library construction and induction

Yeast scFv libraries were constructed by overlap extension PCR using synthetic DNA oligonucleotides. For each sub-library, selected CDR residues were diversified by incorporating NNM codons at the designated positions, where “N” denotes any nucleotide and “M” denotes adenine or cytosine. The V_H_ and V_L_ domains were joined by a (G_4_S)_4_ linker to form the scFv coding sequence. Following PCR assembly, 6 μg of the resulting scFv library DNA and 2 μg of linearized pCTCON2 vector (New England Biolabs) containing a TRP selectable marker were co-transformed into *S. cerevisiae* EBY100 using an electroporation-based method (MicroPulser, Bio-Rad) as previously described.

Transformed yeast were cultured in Trp-deficient SD-CAA medium at 30 °C, harvested, and resuspended at 3 × 10^6^ cells/mL in galactose-containing SG-CAA medium to induce surface display. Cultures were incubated for 20 h at 20 °C to express Aga2p–scFv fusions with a C-terminal Myc tag prior to selection.

### Yeast surface display library construction and screening

Antibody variable fragment libraries were displayed on the surface of *Saccharomyces cerevisiae* EBY100 strain and screened against KRAS(G12D)_10_/HLA-C*08:02/β2m using sequential MACS and FACS. For all selections, cells were washed and resuspended in PBSF buffer (1% BSA in PBS, pH 7.4, 0.22 μm filtered). MACS enrichment was performed with approximately 1 × 10^9^ induced yeast cells, and FACS rounds were performed with 1 × 10^7^ cells.

Biotinylated pMHC I complexes were used for all selection steps. For tetramer-based selections, pMHC I monomers were incrementally added to streptavidin-phycoerythrin (SA-PE) at a defined molar ratio, with the pMHC I in slight molar excess over SA-PE to ensure full tetramerization. For monomer-based selections, biotinylated pMHC I was first incubated with yeast, followed by two washes to remove unbound antigen, after which SA-PE was added to label bound complexes without further oligomerization.

The first library was constructed from nine designed Fv variants, in which three to four residues within the CDR-3 loops were randomized to generate diversity. In particular, for the antibody clone that yielded a binding hit, the initial CDR loop diversification targeted one residue in CDR-H3 and two residues in CDR-L3. Yeast libraries were first enriched by MACS with 100 nM tetramerized target pMHC I, followed by three rounds of FACS positive selection. Upon detection of a distinct binding population, negative selection against SARS_pep_/HLA-C*08:02 was incorporated before positive sorting with the target complex.

A second library, based on the C8K10D framework, incorporated simultaneous diversification of four residues in CDR-H2 and two in CDR-H3. Screening was performed as above but extended to four FACS rounds, with later rounds predominantly using monomeric pMHC I to increase selection stringency.

The third library, derived from C8K10D5, randomized five residues in CDR-L2 and integrated a mammalian cell co-culture step for negative selection. In this approach, yeast were incubated with K562(HLA-C*08:02^+^/TAP^-^/mScarlet-H^+^) cells pulsed with SARS_pep_, and non-binding yeast were recovered for subsequent positive selection with the target monomer. Later rounds incorporated dual-color competition assays in which SARS_pep_/HLA-C*08:02 and KRAS(G12D)_10_/HLA-C*08:02 were labeled with distinct fluorophores, enabling isolation of clones with exclusive binding to the target. The final enrichment steps used <1 nM monomeric target pMHC I to preferentially recover high-affinity binders.

### Antibody production

DNA fragments encoding the V_H_ and V_L_ domains of C8K10D and its derivatives were synthesized (Twist Bioscience) and cloned into a pTT5-derived mammalian expression vector (Addgene).

For Fab production, the V_H_ domain was subcloned into a vector containing the C_H1_ domain with a C-terminal hexahistidine tag, and the VL domain into a vector containing the C_L_ domain. Plasmids encoding the heavy- and light-chain constructs were co-transfected into CHO-S cells (Gibco) at a density of 6 × 10^6^ cells/mL using ExpiFectamine CHO (Gibco), and cultured in ExpiCHO Expression Medium (Gibco) for 10 days. Culture supernatants were clarified by centrifugation at 12,000 × g for 1 h at 4 °C, diluted 1:1 with PBS, and purified by immobilized metal affinity chromatography using HisPur Ni-NTA resin (Thermo Scientific), followed by size-exclusion chromatography on a HiLoad 26/60 Superdex 75 column (Cytiva) equilibrated in PBS.

For full-length IgG production, the VH domain was subcloned into a vector containing the C_H1_–C_H2_–C_H3_ domains, and co-transfected together with the same light chain construct used for Fab expression. Transfection, culture, and clarification steps were identical to those for Fab production. For affinity purification, clarified supernatants were applied to an open column packed with MabSelect^TM^ Sure resin (Cytiva), washed with PBS, and eluted with 0.1 M glycine (pH 3.0). The eluate was immediately neutralized with 1 M Tris-HCl (pH 8.5) and further purified by size-exclusion chromatography using a HiLoad 26/60 Superdex 200 column (Cytiva) equilibrated in PBS.

### Production of peptide–MHC class I complexes

Peptide–MHC I complexes were produced by in vitro protein refolding as previously described. For HLA-C*08:02, KRAS(G12D)_10_ peptide (GVDGVGKSAL) as the target and a non-target control peptide (SARS_pep_; ITDAVDCAL) were used. For HLA-A02:01, the p53(R175H) peptide (HMTEVVRHC) was used as a control antigen. In brief, denatured HLA heavy chain and β2m were refolded in the presence of the corresponding synthetic peptide in a refolding buffer containing 20 mM Tris-HCl (pH 8.0), 400 mM L-arginine, 2 mM EDTA, 0.5 mM oxidized glutathione, 5 mM reduced glutathione, and 0.2 mM phenylmethylsulfonyl fluoride (PMSF). The reaction was incubated at 18 °C for 48 h, concentrated to 7.5 mL, and buffer-exchanged using a PD-10 column equilibrated with 30 mM NaCl, 20 mM Tris-HCl (pH 8.0). The resulting complexes were further purified by size exclusion chromatography using a HiPrep™ 26/60 Sephacryl® S-300 HR column (Cytiva), immobilized metal affinity chromatography (Ni-NTA), and anion exchange chromatography on a Capto™ HiRes Q 5/50 column (Cytiva).

### Production of biotinylated peptide–MHC class I complexes

For site-specific biotinylation, an Avi-tag sequence was appended to the C-terminus of the HLA heavy chain by molecular cloning. Refolding was performed as above, except that the PD-10 buffer exchange step used biotinylation buffer containing 10 mM Tris-HCl (pH 7.5), 200 mM NaCl, and 5 mM MgCl_2_. The refolded complex was supplemented with 5 mM ATP, 400 µM biotin, leupeptin and pepstatin, 200 µM PMSF, and 60 nM BirA enzyme, and incubated at 4 °C overnight. The biotinylated pMHC I was then purified by immobilized metal affinity chromatography (Ni-NTA), size exclusion chromatography using a HiPrep™ 26/60 Sephacryl® S-300 HR column (Cytiva), and anion exchange chromatography on a Capto™ HiRes Q 5/50 column (Cytiva).

### Biolayer interferometry

Binding kinetics were measured using an Octet R8 system (Sartorius). Biotinylated pMHC I complexes were immobilized onto streptavidin (SA) biosensor tips (Sartorius) at a concentration of 2.5 nM in Kinetics Buffer (Sartorius). For all experiments, double reference subtraction was performed against both an empty ligand control and an empty analyte control to account for non-specific binding to the sensor tip. Association and dissociation times were adjusted according to the observed target binding affinity, typically ranging from 120 s to 480 s for association and dissociation. Binding data were analyzed using the Octet BLI Analysis software version 12.2.2.4 (Sartorius).

### Generation of stable cell line

Stable 721.221 and K562 (HLA C*08:02^+^/TAP^-^) cell lines were generated by sequential transduction using lentiviral knock-in and lentivirus mediated CRISPR/cas9 knock-out system, as previously described (RMF paper). A second-generation CAR construct was engineered by linking scFVs of the C8K10D5 variants via a Gly-Ser (GS) linker to the CD8α hinge and transmembrane domains, followed by the intracellular signaling domains of 4-1BB and CD3ζ. A truncated LNGFR (ΔLNGFR, CD271) served as a selection marker. The CAR expression cassette was inserted into the pLV-EF1a-IRES-Puro lentiviral vector, replacing the IRES-Puro element with p2a-ΔLNGFR. The lentiviral transduction of CAR-expressing cells was performed as previously described (40).

### Mammalian Cell Culture

The human B-lymphoblastoid cell line 721.221 (ATCC CRL-1855) cells were maintained in RPMI1640 (Gibco, 72400047) supplemented with 10% Fetal Bovine Serum, (FBS; Gibco, 12483020). The human erythromyeloid leukemia cell line K562 (ATCC CRL-243) was maintained in IMDM medium (Gibco, 12440053) supplemented with 10% FBS. All cells were maintained at 37 °C in a 5% CO_2_ humidified incubator.

### Peptide pulsing

721.221 (HLA-C*08:02^+^/TAP^-^) and K562 (HLA-C*08:02^+^/TAP^-^) cells were washed with serum-free RPMI 1640 or IMDM, respectively. The cells were transferred to low attachment plates (SPL Life Sciences, 32006) with the concentration of 1-2 x 10^6^ cells/mL. For peptide-pulsing condition, 10 µg/mL human β2m with each of 10µM KRAS(G12D)_10_, SARS_pep_, KRAS(G12V), KRAS(G12R), KRAS(G12C), KRAS WT, or peptide-substituted KRAS(G12D)_10_ peptides were added to each well. For non-pulsed condition, only 10 µg/mL human β2m was added. Then, the plates were incubated for 20-24 hours at 26 °C in a humidified incubator with 5% CO_2_ prior to subsequent functional assays.

### Cellular binding assay

Pulsed cells were washed once with the ice-cold PBSF buffer and resuspended at 2–5 × 10^5^ cells per 100 µL. Cells were incubated for 1 hour with 50 nM of each Fab clone or full antibody for primary staining, followed by 1 hour with 50 nM Fluorescein isothiocyanate (FITC) Anti-6X His tag® antibody (Abcam, ab1206) or Protein A, Alexa Fluor™ 488 conjugate (Invitrogen, P11047) for secondary staining at 4 °C in the dark. After each staining step, cells were washed twice with the PBSF buffer. The cells were then resuspended in 200 µL of the PBSF buffer and analyzed on a CytoFLEX SRT cell sorter (Beckman Coulter). Binding quantification was measured by mean fluorescence intensity within the live-cell gating.

### Confocal imaging

Pulsed K562 (HLA-C*08:02^+^/TAP^-^/mScarlet-H^+^) cells were resuspended at 2 x 10^5^ cells in 200 µL PBSF buffer and Yeast cells were suspended at 20 x 10^5^ cells in 200 µL. The cells were then co-incubated at a ratio of 1:10 for 10 minutes. Following incubation, all cells were added 1600 µL of PBSF buffer and plated on 35mm glass-bottom confocal dishes (SPL, 200350) for imaging. Live-cell imaging was performed using a Nikon A1HD25microscope (Nikon Instruments) mounted on a Nikon Eclipse Ti2 inverted microscope, equipped with a PLAN APO lambda (x40, 0.95 N.A., Nikon Instruments). The images were acquired using digital-zooming Nikon imaging software (NIS-element AR 5.30.00).

### Structure determination

Optimized crystallization conditions for C8K10D5-1 Fab/KRAS(G12D)/HLA-C*08:02/β2m were established by hanging-drop vapor diffusion at 20 °C, using 2 μL drops prepared by mixing equal volumes of protein and reservoir solutions. C8K10D5-1 Fab/KRAS(G12D)_10_/HLA-C*08:02/β2m crystals were obtained using the protein sample (16 mg/mL) and a reservoir solution composed of 1200 mM NaH_2_PO_4_, 200 mM (NH_4_)_2_SO_4_, 200mM Li_2_SO_4_ and 100 mM CAPS (pH 10.5).

For cryoprotection, the crystals were transferred into the reservoir solution supplemented with 15.0 % (v/v) glycerol and subsequently flash-frozen in liquid nitrogen. X-ray diffraction data were collected at beamline 5C of Pohang Accelerator Laboratory, and processed with HKL2000. The model was refined using COOT and PHENIX.

### Production of CAR-T cells

Primary human peripheral blood mononuclear cells (PBMCs) were purchased from Miltenyi Biotec. PBMCs were activated with Dynabeads™ Human T-Activator CD3/CD28 (Thermo Fisher Scientific) at a 1:1 bead-to-cell ratio and maintained in RPMI-1640 medium (Gibco) supplemented with 10% FBS and 300 IU/mL IL-2.

Activated T cells were transduced with lentiviral vectors encoding second-generation CAR constructs incorporating scFvs derived from C8K10D5 variants (C8K10D5-1, -18, -18-DE, and -18-DQE). Transduced cells were expanded for 7–10 days in the presence of IL-2, and CAR expression was confirmed by flow cytometry using anti-LNGFR (CD271) staining.

### Cytotoxicity assay of CAR-T cells

Cytotoxic activity was measured using K562(HLA-C08:02^+^/TAP^-^) and 721.221(HLA-C08:02^+^/TAP^-^) target cells pulsed with 10 μM KRAS(G12D)_10_ peptide, a control peptide (SARS_pep_), or left unpulsed. Effector CAR-T cells were co-cultured with target cells at effector-to-target (E:T) ratios of 1:1 and 1:5 in 96-well round-bottom plates for 4 h at 37 °C in 5% CO_2_. Target cell lysis was quantified using a calcein-AM release assay (Thermo Fisher Scientific). Briefly, target cells were labeled with 5 μM calcein-AM for an hour at 37 °C, washed, and co-cultured with CAR-T cells. Fluorescence in the supernatant was measured with a microplate reader at 494/535 nm. Spontaneous release was defined as fluorescence from calcein-AM–labeled target cells incubated without effector cells (target only), and maximum release was determined by treating labeled target cells with lysis solution. Specific lysis (%) was calculated as:

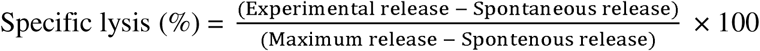

All experiments were performed in triplicate and analyzed using GraphPad Prism.

## Acknowledgments

This study utilized the Beamline 5C at the Pohang Accelerator Laboratory, Republic of Korea.

## Author contributions

Conceptualization: B.-S.J. and B.-H.O.

Methodology and Investigation: B.-S.J., S.A., H.L., J.L., S.S., T.-S.O., J.Y.J., K.R., H.K., E.G.L., W.J., B.M.K., J.J, M.-J.A. and D.K.K.

Validation: B.-S.J., H.L., J.L., and S.A.

Formal analysis: B.-S.J., S.A., and S.L.

Data curation: B.-S.J., S.A., S.L., and B.-H.O.

Writing—original draft: B.-H.O., B.-S.J., S.A., and S.L.

Writing—review and editing: All authors

Visualization: B.-S.J. and S.A.

Supervision: B.-H.O. and B.-S.J.

Project administration: B.-H.O. and B.-S.J.

Funding acquisition: B.-H.O.

Resources: B.-H.O.

## Data and materials availability

All data needed to evaluate the conclusions in the paper are present in the paper and/or the Supplementary Materials.The coordinates of the C8K10D5-1 Fab/KRAS(G12D)_10_/HLA-C*08:02/β2m structure have been deposited in the Protein Data Bank (PDB ID: 9WJ6) with the conditions of immediate release upon publication.

## Supplementary figure legend

**Fig. S1. YSD screening of the initial design with randomized CDR-3 loops.**

Yeast surface display screening of the first scFv library. Nine designed Fv variants were used to construct a CDR-3-focused library in which three to four residues were randomized. For the clone that yielded a binding hit, one CDR-H3 residue and two CDR-L3 residues were diversified. A single clone became enriched after iterative sorting. From the point at which enrichment was detectable, binding to SARS_pep_/HLA-C08:02 was also monitored, and sorting was conducted only for clones retaining specificity to the target pMHC. No cross-reactivity to SARS_pep_/HLA-C*08:02 was detected under these screening conditions (data not shown).

**Fig. S2. YSD screening of C8K10D with targeted CDR-H2 and CDR-H3 randomization.**

Four residues in CDR-H2 and two residues in CDR-H3 were selected for diversification. The library was screened using the same workflow as for the initial design, with four rounds of FACS. Monomeric KRAS(G12D)_10_/HLA-C*08:02 was primarily used for positive selection during this sorting procedure.

**Fig. S3. Affinity measurement of C8K10D3-Fab and C8K10D6-Fab.**

Binding kinetics were determined by BLI. Biotinylated target pMHC was immobilized on SA biosensor tips, and Fabs were tested at an initial concentration of 125 nM followed by serial twofold dilutions. Association and dissociation phases were recorded, and kinetic parameters were obtained by global fitting of the resulting binding curves.

**Fig. S4. Confocal Images of K562(HLA-C*08:02^+^/TAP^-^) cells, MOCK-displaying yeast, and C8K10D5 scFv-displaying yeast.**

Top row: self-incubation controls of each cell type. Middle row: MOCK-displaying yeast (green) co-incubated with K562(HLA-C*08:02^+^/TAP^-^) cells (red) pulsed with SARS_pep_ peptide, KRAS(G12D)_10_ peptide or left unpulsed. No binding was observed in any condition. Bottom Row: C8K10D5 scFv–displaying yeast (green) co-incubated with K562(HLA-C*08:02^+^/TAP^-^) cells (Red) pulsed with SARS_pep_ peptide, KRAS(G12D)_10_ peptide or left unpulsed. Specific binding was observed solely in the KRAS(G12D)_10_ peptide-pulsed condition.

**Fig. S5. YSD screening of the C8K10D5 sub-library using mammalian cell co-culture.**

Five residues within CDR-L2 were randomized to generate the sub-library. The library was first subjected to MACS for enrichment. The first round of FACS involved positive selection using the target pMHC protein. In the second and third rounds, yeast clones were co-cultured with mammalian cells presenting a non-target pMHC complex (K562 (HLA-C*08:02^+^/TAP^-^/mScarlet-H^+^) pulsed with SARS_pep_) for negative selection, followed by positive sorting for binding to the target pMHC protein. The fourth round involved positive selection for target pMHC I monomer binding in the presence of unpulsed K562 cells. In the fifth round, SARS_pep_/HLA-C08:02 and KRAS(G12D)_10_/HLA-C*08:02 were differentially labeled with distinct fluorophores, and only populations binding selectively to the target were sorted. The sixth and seventh rounds employed sub-nanomolar concentrations (<1 nM) of target pMHC I monomer to enrich high-affinity binders.

**Fig. S6.** Flow cytometric analysis of yeast–mammalian co-culture sorting outcome.

Binding of the C8K10D5 sub-library to K562(HLA-C*08:02^+^/TAP^-^) cells was evaluated before and after the mammalian co-culture sorting process. Before sorting, a substantial fraction of yeast bound to non-pulsed K562 cells. Following sorting, this non-specific binding population was markedly reduced. In contrast, binding to KRAS(G12D)_10_-pulsed K562 cells was substantially enriched in the sorted population.

**Fig. S7. Analysis of peptide specificity using alanine-scanning substitutions at each position of the 10-mer KRAS(G12D) peptide in two cell lines.**

721.221(HLA-C*08:02^+^/TAP^-^) cells (top panels) and K562(HLA-C*08:02^+^/TAP^-^) cells (bottom panels) were pulsed with the alanine-substituted peptides and subsequently analyzed by flow cytometry. Binding was quantified by measuring the mean fluorescence intensity within live cell gates using the FITC channel. Dashed lines represent the average Mean-FITC values from non-pulsed control samples stained with C8K10D5-1 Fab and C8K10D5-18 Fab. Both Fab clones showed loss of binding to specific alanine-substituted peptide variants. No binding was detected against the wild-type peptide or other KRAS G12 mutations.

**Fig. S8. Affinity measurement of C8K10D5-18-DE Fab and C8K10D5-18-DQE Fab.**

Binding kinetics were determined by BLI. Biotinylated target pMHC was immobilized on SA biosensor tips, and Fabs were tested at an initial concentration of 32 nM followed by serial twofold dilutions. Association and dissociation phases were recorded, and kinetic parameters were obtained by global fitting of the resulting binding curves.

**Fig. S9. Specificity profiling of Fab binding on antigen-presenting cell lines.**

K562(HLA-C*08:02^+^/TAP^-^) (Left) and 721.221(HLA-C*08:02^+^/TAP^-^) (right) cells were pulsed with the KRAS(G12D)_10_ peptide, a control SARS_pep_ peptide, or left unpulsed. Fab binding was assessed by flow cytometry and quantified as mean fluorescence intensity within live cell gates in the FITC channel. Dashed lines represent the average Mean-FITC values of non-pulsed control samples stained with each Fab. C8K10D5-1, C8K10D5-18, C8K10D5-18-DE, and C8K10D5-18-DQE Fabs demonstrated specific binding to cells pulsed with the KRAS(G12D)_10_ peptide.

**Fig. S10. CAR expression and peptide-dependent cytotoxicity of C8K10D5 variant–derived CAR-T cells.**

**A.** Representative flow-cytometry plots (SSC-A vs dLNGFR) showing CAR expression in primary T cells transduced with the indicated constructs (CAR-C8K10D5-1, -18, -18-DE, -18-DQE). Numbers denote the percentage of dLNGFR^+^ cells. NTD, non-transduced T-cell control.

**B.** Peptide-dependent cytotoxicity of C8K10D5 variant–derived CAR-T cells. K562(HLA-C*08:02^+^/TAP^-^) cells and 721.221(HLA-C*08:02^+^/TAP^-^) target cells were pulsed with KRAS(G12D)_10_ or SARS_pep_ peptides or left unpulsed, and co-cultured with T cells expressing CARs derived from C8K10D5 variants (C8K10D5-1, -18, -18-DE, -18-DQE) at an effector-to-target ratio of 1:1 for 4 h. Cytotoxicity was measured by calcein-AM release assay. Data are represented mean ± SD of triplicates; statistical analyses were performed using GraphPad Prism.

