## Supplementary figures for "An integrated in silico-in vitro workflow for discovering high-affinity, selective antibodies to the KRAS(G12D)-MHC I complex"

### Slide 1
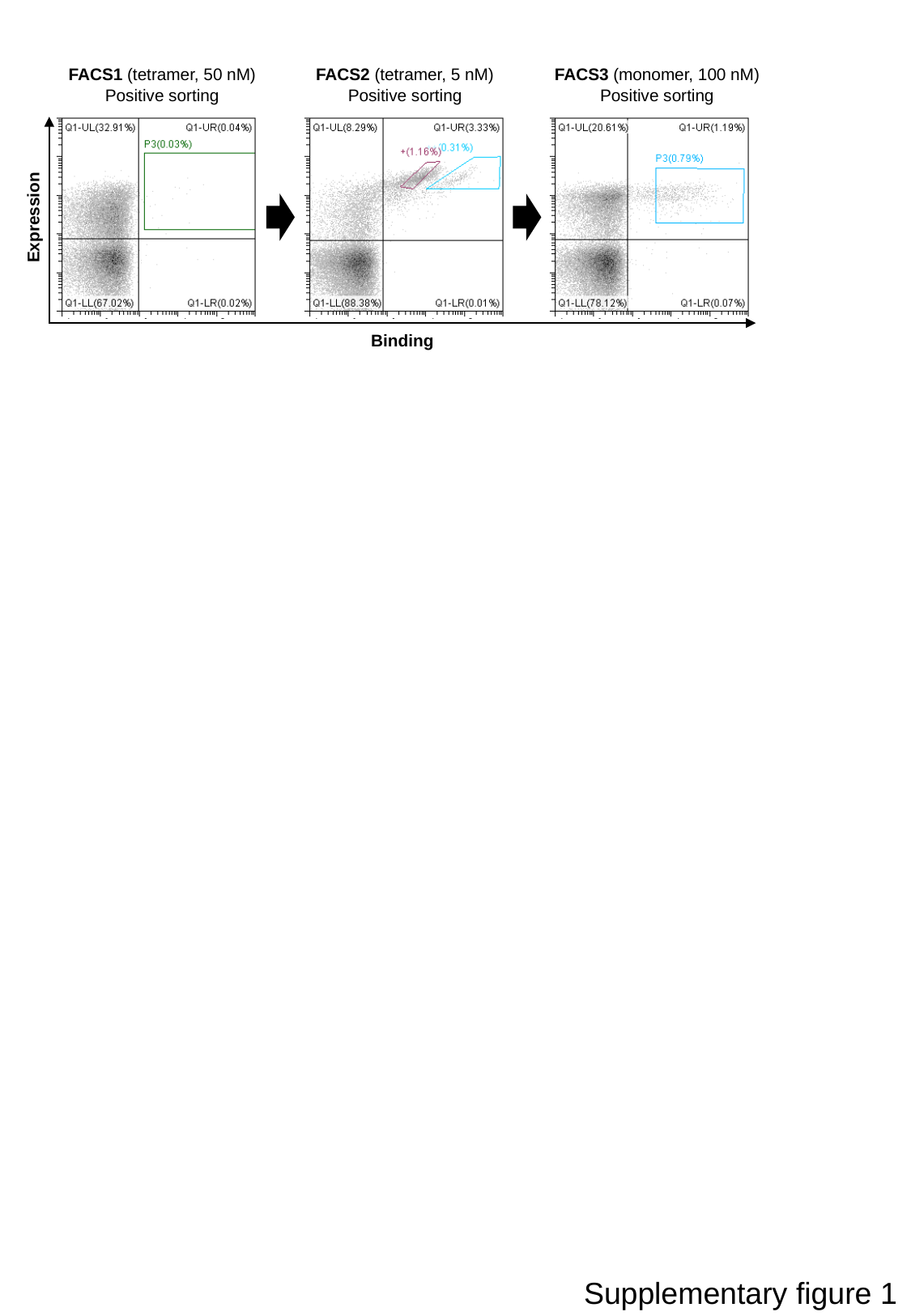

FACS1 (tetramer, 50 nM)
Positive sorting
FACS2 (tetramer, 5 nM)
Positive sorting
FACS3 (monomer, 100 nM)
Positive sorting
Expression
Binding
Supplementary figure 1

### Slide 2
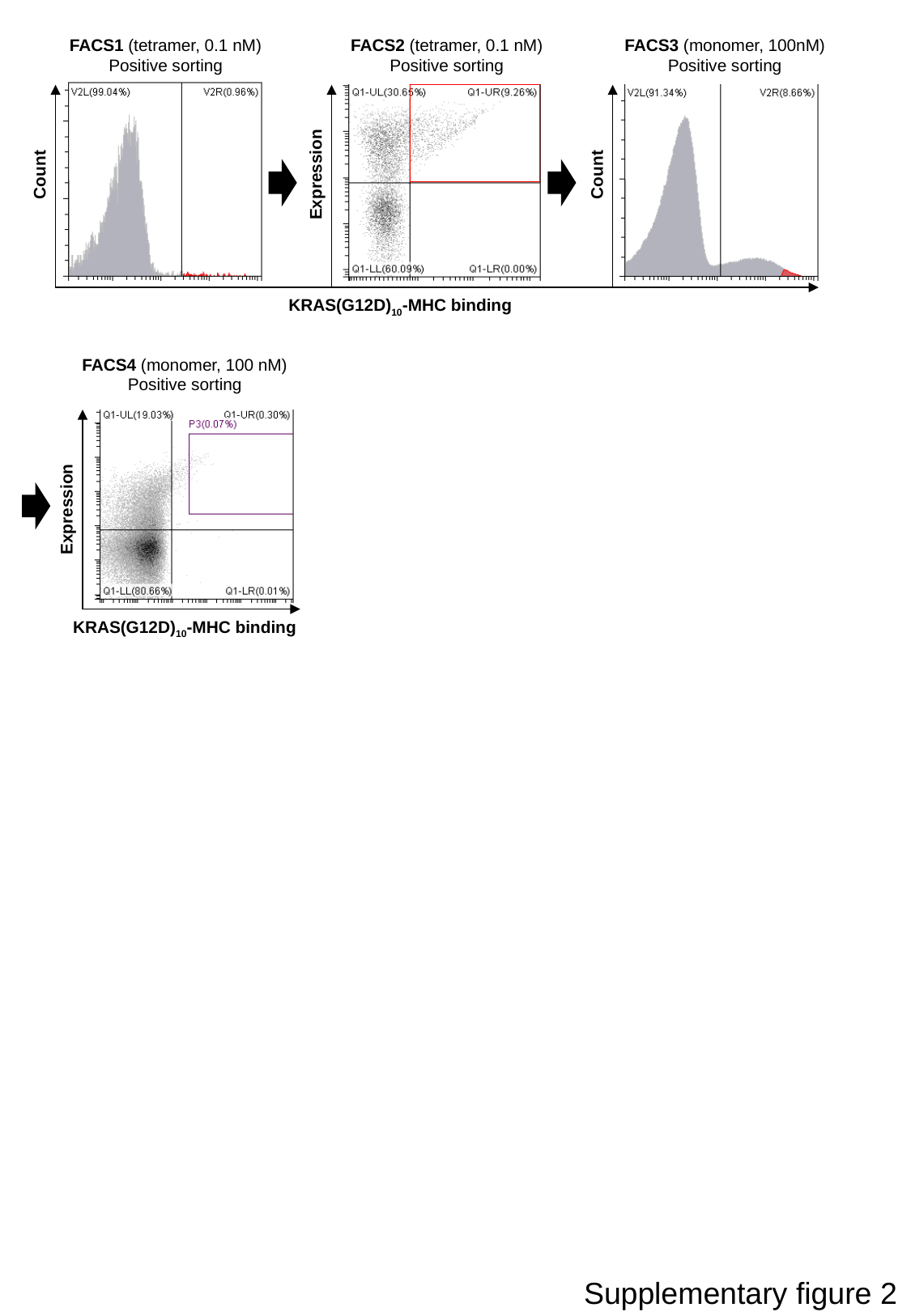

FACS1 (tetramer, 0.1 nM)
Positive sorting
FACS2 (tetramer, 0.1 nM)
Positive sorting
FACS3 (monomer, 100nM)
Positive sorting
Count
Expression
Count
KRAS(G12D)_10_-MHC binding
FACS4 (monomer, 100 nM)
Positive sorting
Expression
KRAS(G12D)_10_-MHC binding
Supplementary figure 2

### Slide 3
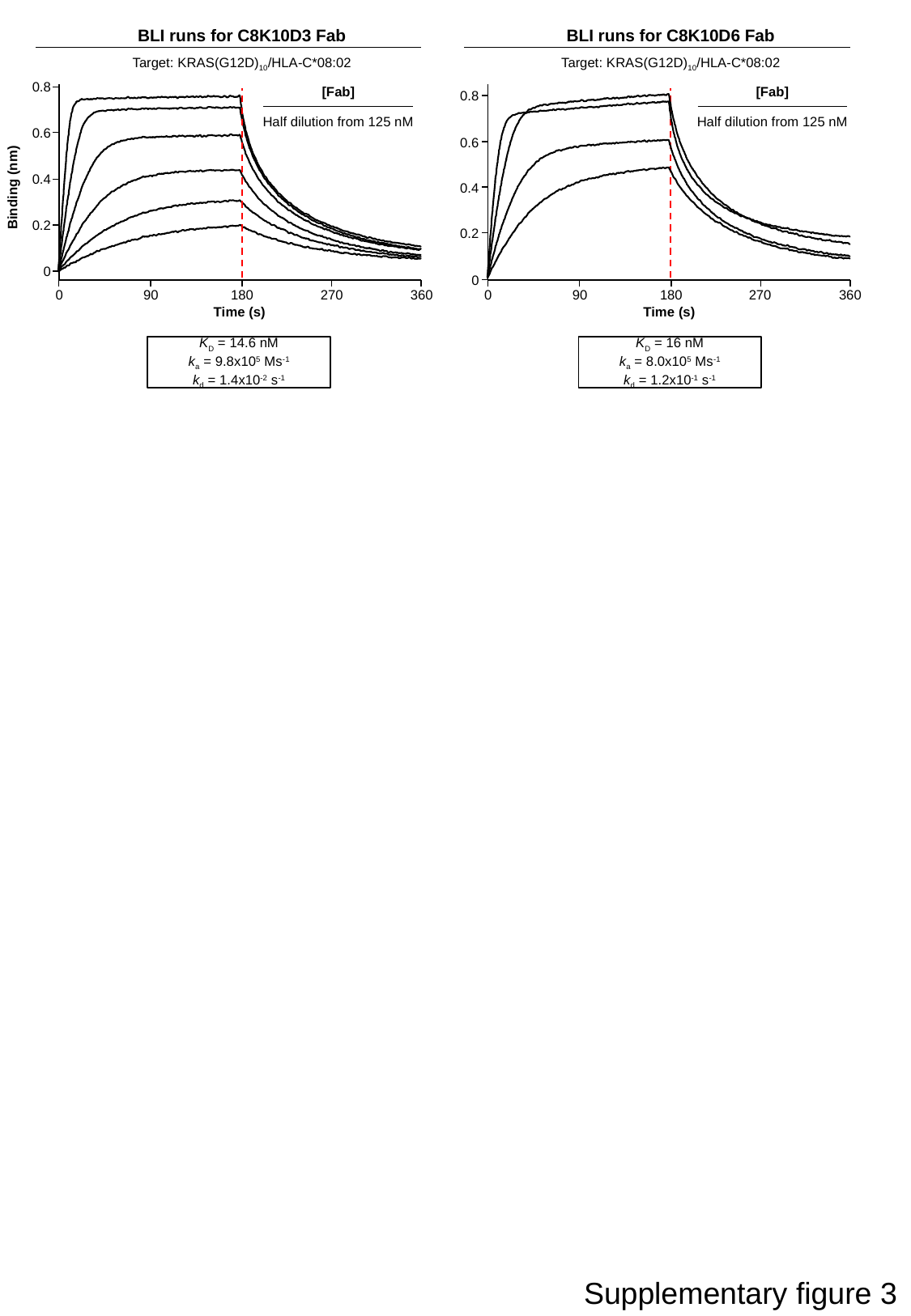

BLI runs for C8K10D3 Fab
Target: KRAS(G12D)_10_/HLA-C*08:02
BLI runs for C8K10D6 Fab
Target: KRAS(G12D)_10_/HLA-C*08:02
0.8
[Fab]
Half dilution from 125 nM
[Fab]
Half dilution from 125 nM
0.8
0.6
0.6
0.4
Binding (nm)
0.4
0.2
0.2
0
0
0
90
180
270
360
0
90
180
270
360
Time (s)
Time (s)
K_D_ = 14.6 nM
k_a_ = 9.8x10^5^ Ms^-1^
k_d_ = 1.4x10^-2^ s^-1^
K_D_ = 16 nM
k_a_ = 8.0x10^5^ Ms^-1^
k_d_ = 1.2x10^-1^ s^-1^
Supplementary figure 3

### Slide 4
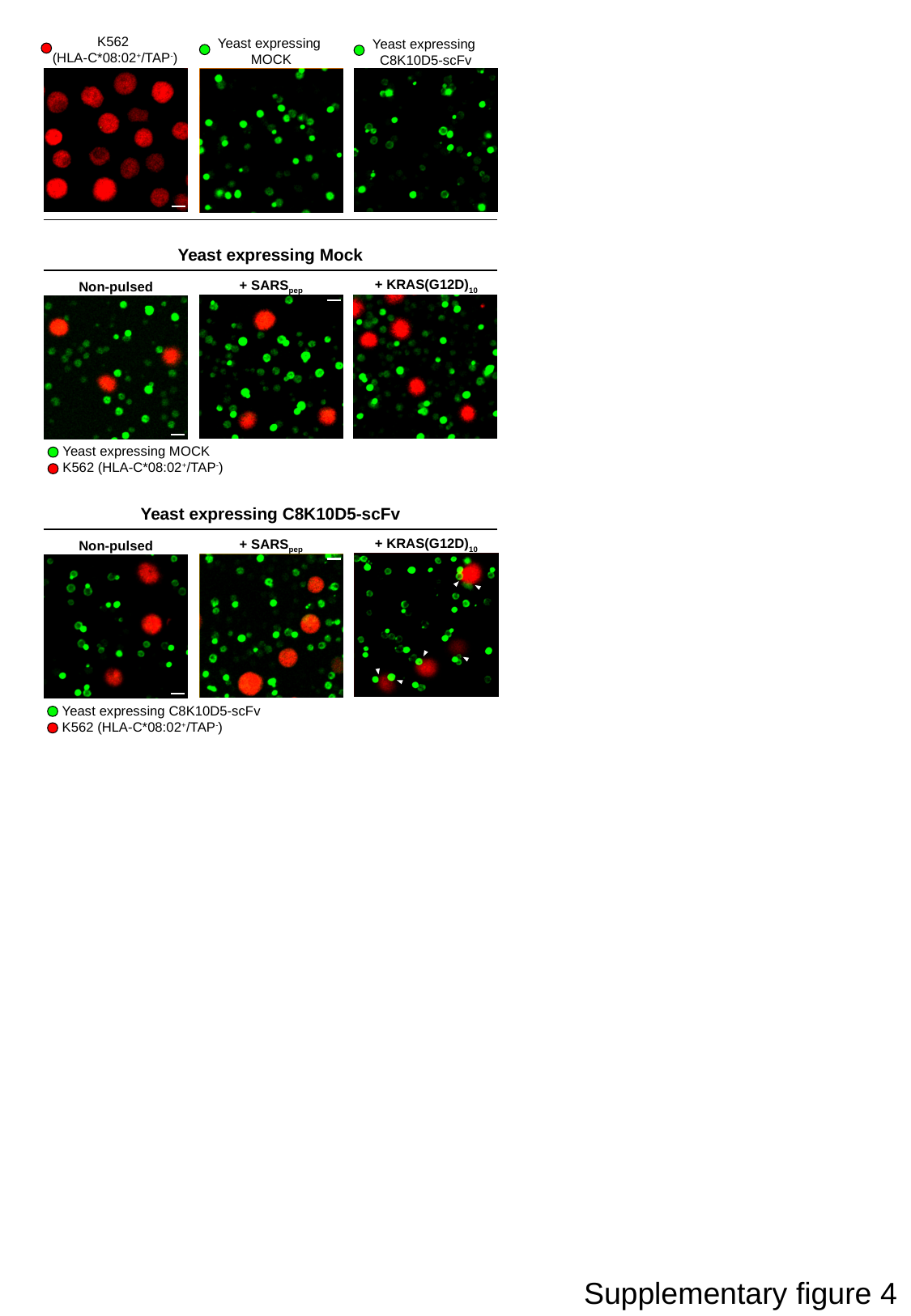

K562
(HLA-C*08:02^+^/TAP^-^)
Yeast expressing
MOCK
Yeast expressing
C8K10D5-scFv
Yeast expressing Mock
+ KRAS(G12D)_10_
+ SARS_pep_
Non-pulsed
Yeast expressing MOCK
K562 (HLA-C*08:02^+^/TAP^-^)
Yeast expressing C8K10D5-scFv
+ KRAS(G12D)_10_
+ SARS_pep_
Non-pulsed
Yeast expressing C8K10D5-scFv
K562 (HLA-C*08:02^+^/TAP^-^)
Supplementary figure 4

### Slide 5
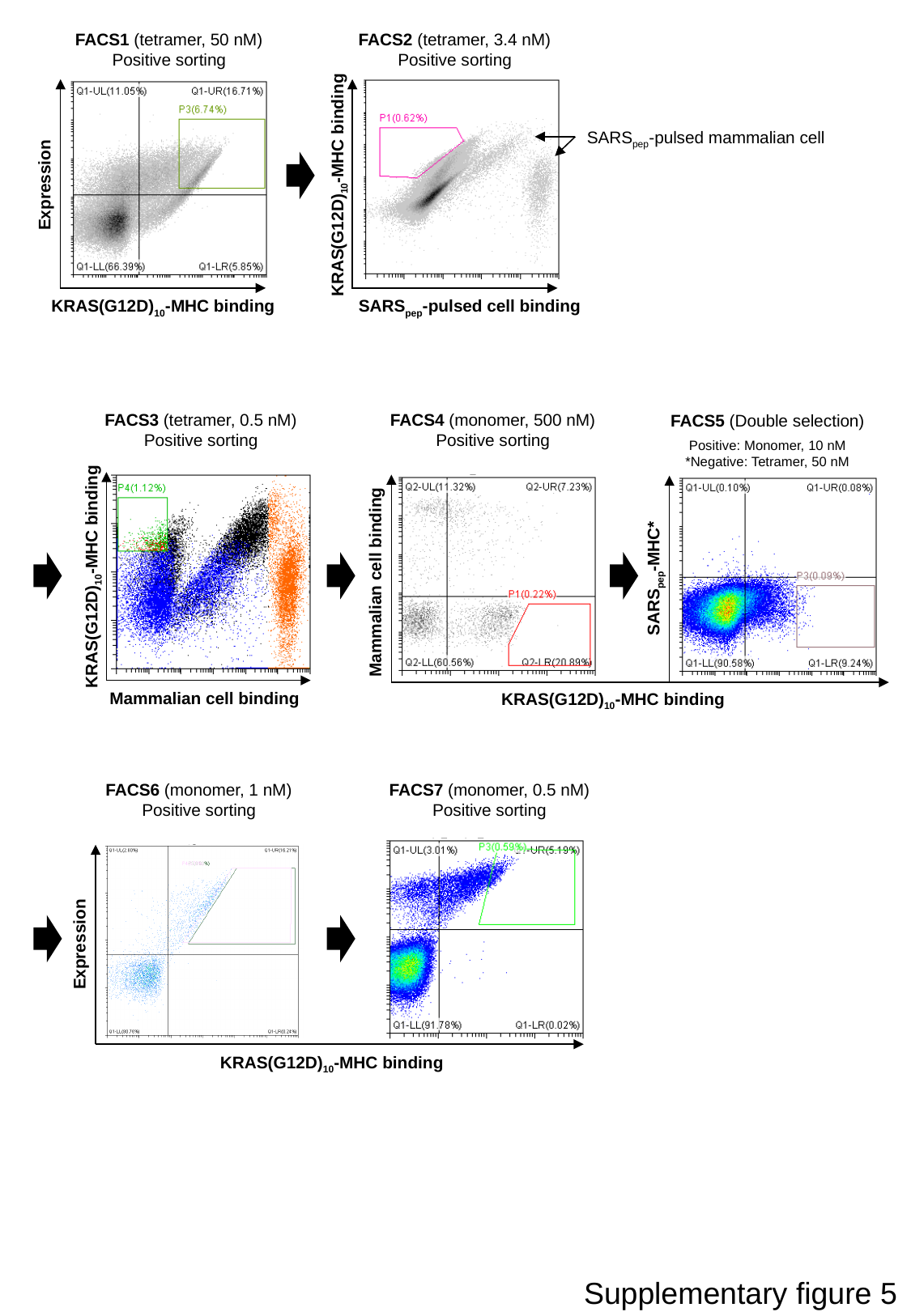

FACS1 (tetramer, 50 nM)
Positive sorting
FACS2 (tetramer, 3.4 nM)
Positive sorting
SARS_pep_-pulsed mammalian cell
KRAS(G12D)_10_-MHC binding
Expression
KRAS(G12D)_10_-MHC binding
SARS_pep_-pulsed cell binding
FACS3 (tetramer, 0.5 nM)
Positive sorting
FACS4 (monomer, 500 nM)
Positive sorting
FACS5 (Double selection)
Positive: Monomer, 10 nM
*Negative: Tetramer, 50 nM
KRAS(G12D)_10_-MHC binding
SARS_pep_-MHC*
Mammalian cell binding
Mammalian cell binding
KRAS(G12D)_10_-MHC binding
FACS6 (monomer, 1 nM)
Positive sorting
FACS7 (monomer, 0.5 nM)
Positive sorting
Expression
KRAS(G12D)_10_-MHC binding
Supplementary figure 5

### Slide 6
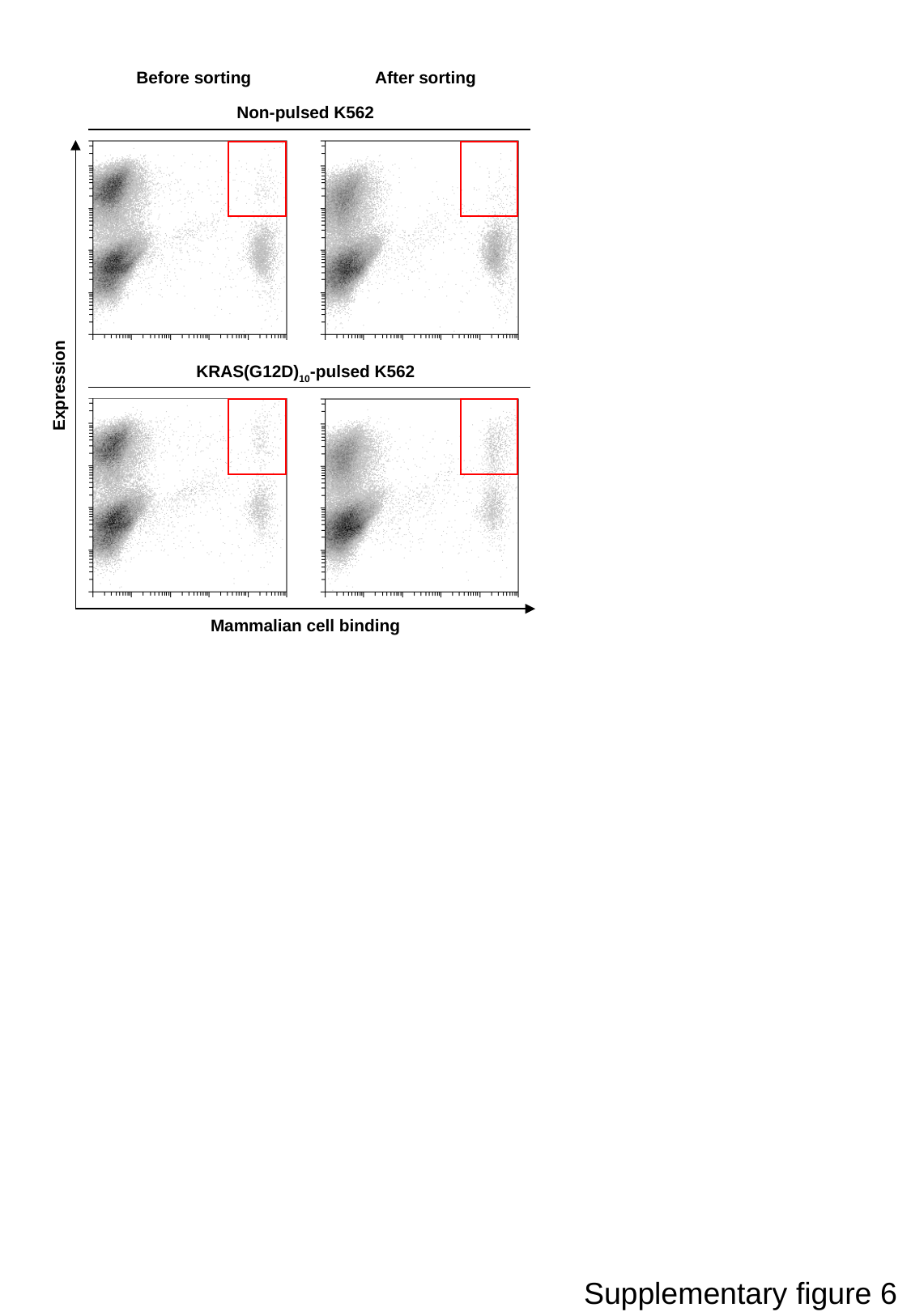

Before sorting
After sorting
Non-pulsed K562
KRAS(G12D)_10_-pulsed K562
Expression
Mammalian cell binding
Supplementary figure 6

### Slide 7
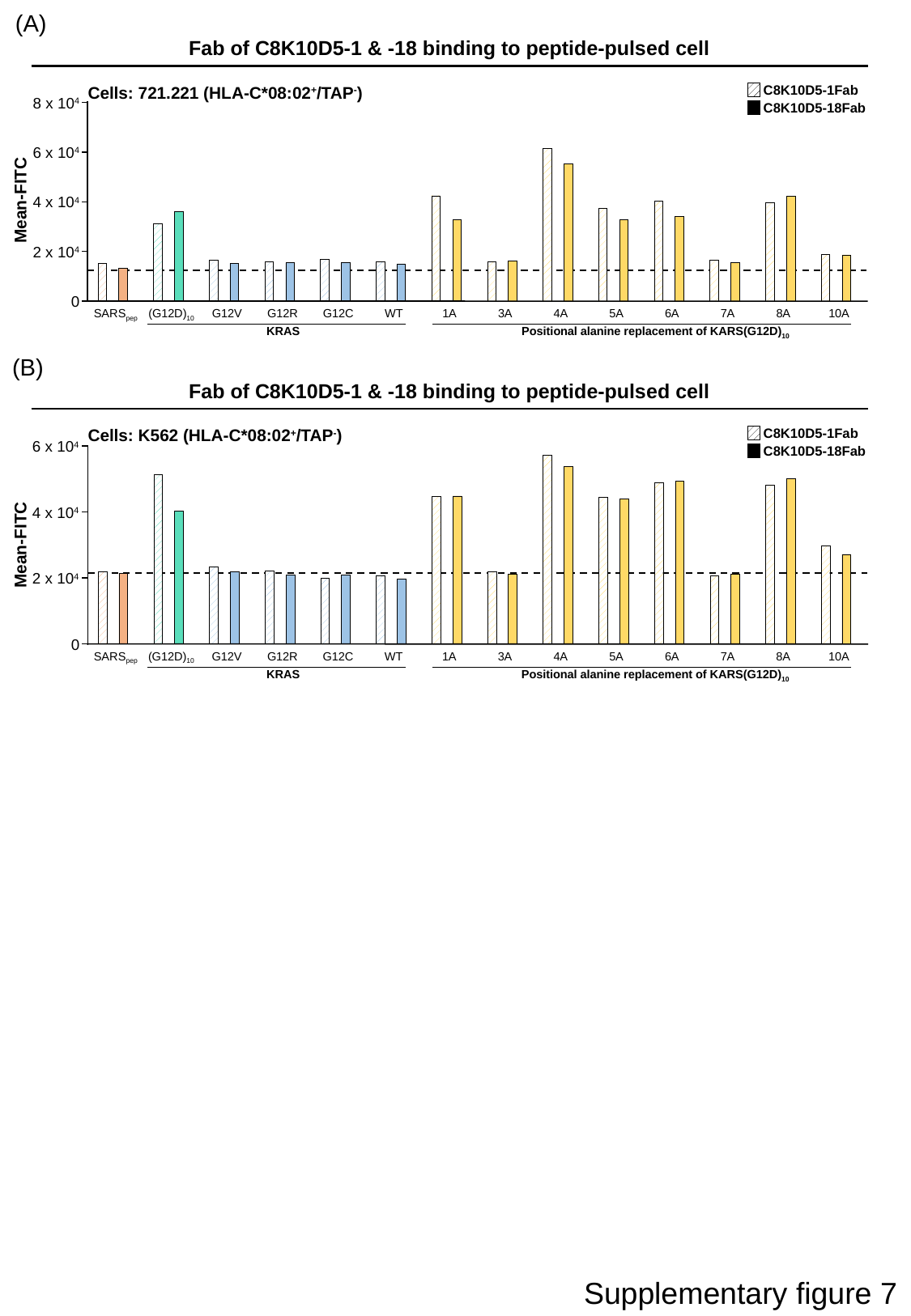

(A)
Fab of C8K10D5-1 & -18 binding to peptide-pulsed cell
C8K10D5-1Fab
C8K10D5-18Fab
Cells: 721.221 (HLA-C*08:02^+^/TAP^-^)
8 x 10^4^
6 x 10^4^
4 x 10^4^
2 x 10^4^
0
SARS_pep_
(G12D)_10_
G12V
G12R
G12C
WT
1A
3A
4A
5A
6A
7A
8A
10A
KRAS
Positional alanine replacement of KARS(G12D)_10_
Mean-FITC
(B)
Fab of C8K10D5-1 & -18 binding to peptide-pulsed cell
C8K10D5-1Fab
C8K10D5-18Fab
Cells: K562 (HLA-C*08:02^+^/TAP^-^)
6 x 10^4^
4 x 10^4^
2 x 10^4^
0
SARS_pep_
(G12D)_10_
G12V
G12R
G12C
WT
1A
3A
4A
5A
6A
7A
8A
10A
KRAS
Positional alanine replacement of KARS(G12D)_10_
Mean-FITC
Supplementary figure 7

### Slide 8
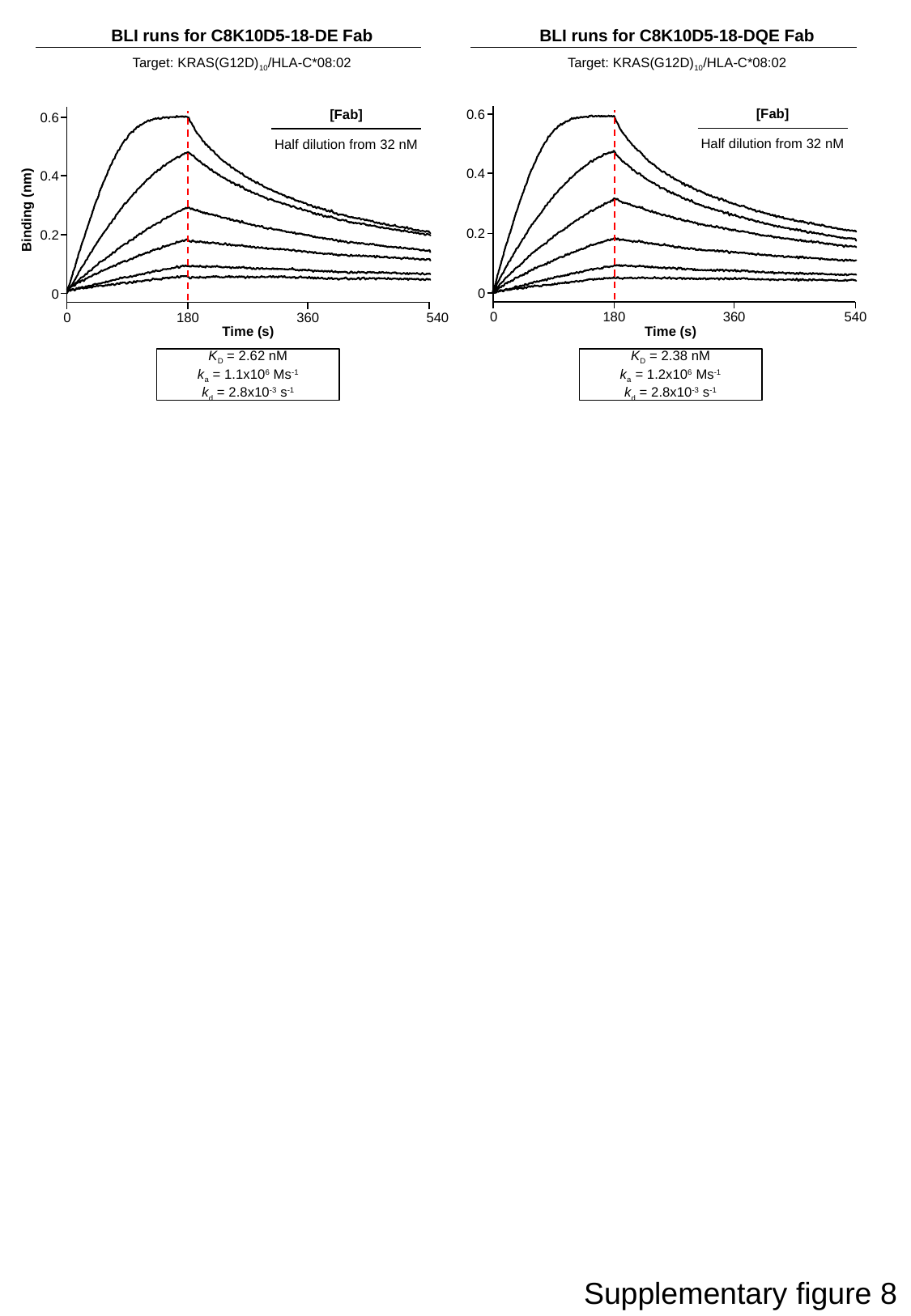

BLI runs for C8K10D5-18-DE Fab
Target: KRAS(G12D)_10_/HLA-C*08:02
BLI runs for C8K10D5-18-DQE Fab
Target: KRAS(G12D)_10_/HLA-C*08:02
[Fab]
Half dilution from 32 nM
0.6
[Fab]
Half dilution from 32 nM
0.6
0.4
0.4
Binding (nm)
0.2
0.2
0
0
0
180
360
540
0
180
360
540
Time (s)
Time (s)
K_D_ = 2.62 nM
k_a_ = 1.1x10^6^ Ms^-1^
k_d_ = 2.8x10^-3^ s^-1^
K_D_ = 2.38 nM
k_a_ = 1.2x10^6^ Ms^-1^
k_d_ = 2.8x10^-3^ s^-1^
Supplementary figure 8

### Slide 9
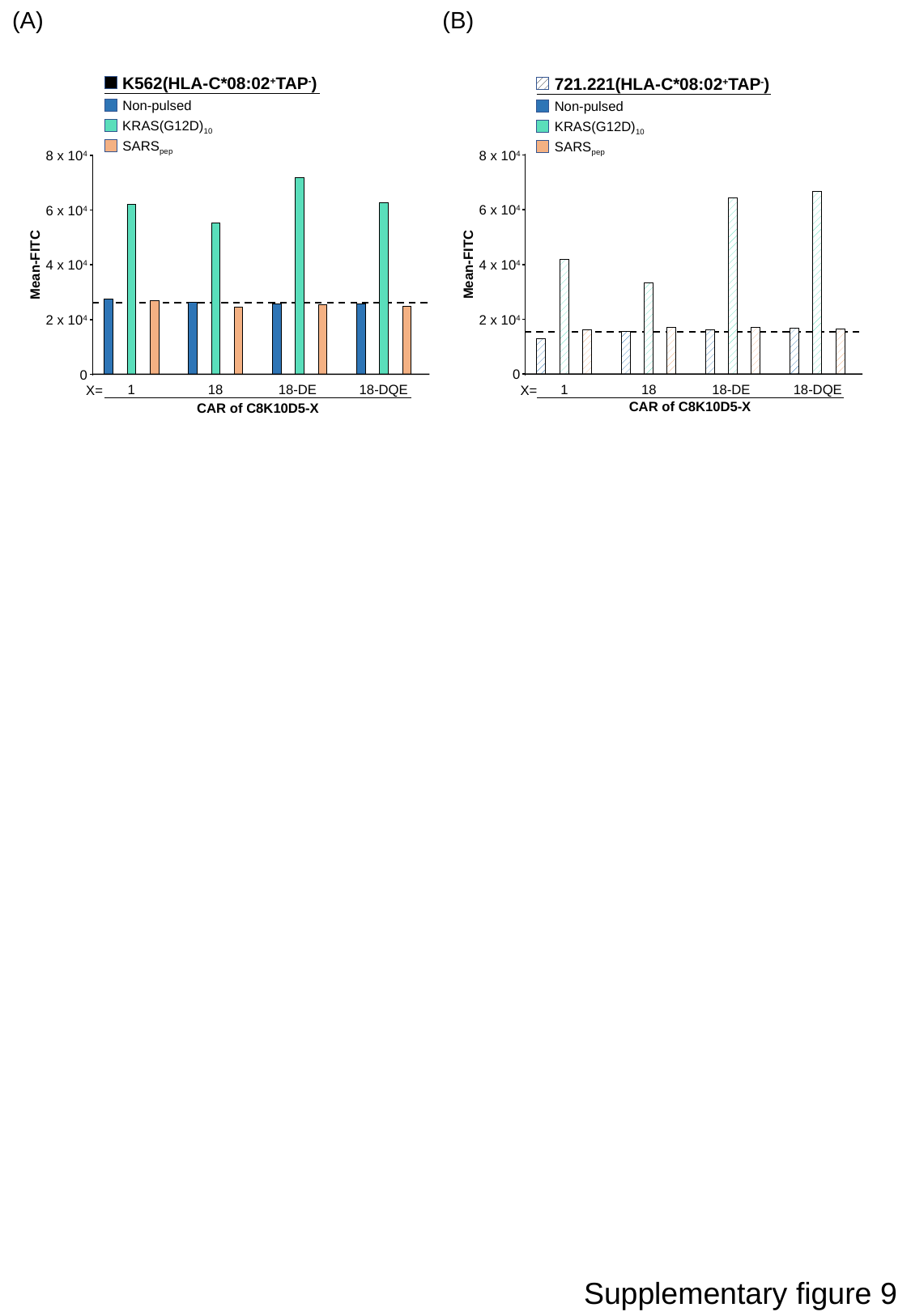

(A)
(B)
K562(HLA-C*08:02^+^TAP^-^)
Non-pulsed
KRAS(G12D)_10_
SARS_pep_
721.221(HLA-C*08:02^+^TAP^-^)
Non-pulsed
KRAS(G12D)_10_
SARS_pep_
 8 x 10^4^
 8 x 10^4^
6 x 10^4^
6 x 10^4^
4 x 10^4^
Mean-FITC
4 x 10^4^
Mean-FITC
2 x 10^4^
2 x 10^4^
0
0
1
18
18-DE
18-DQE
1
18
18-DE
18-DQE
X=
X=
CAR of C8K10D5-X
CAR of C8K10D5-X
Supplementary figure 9

### Slide 10
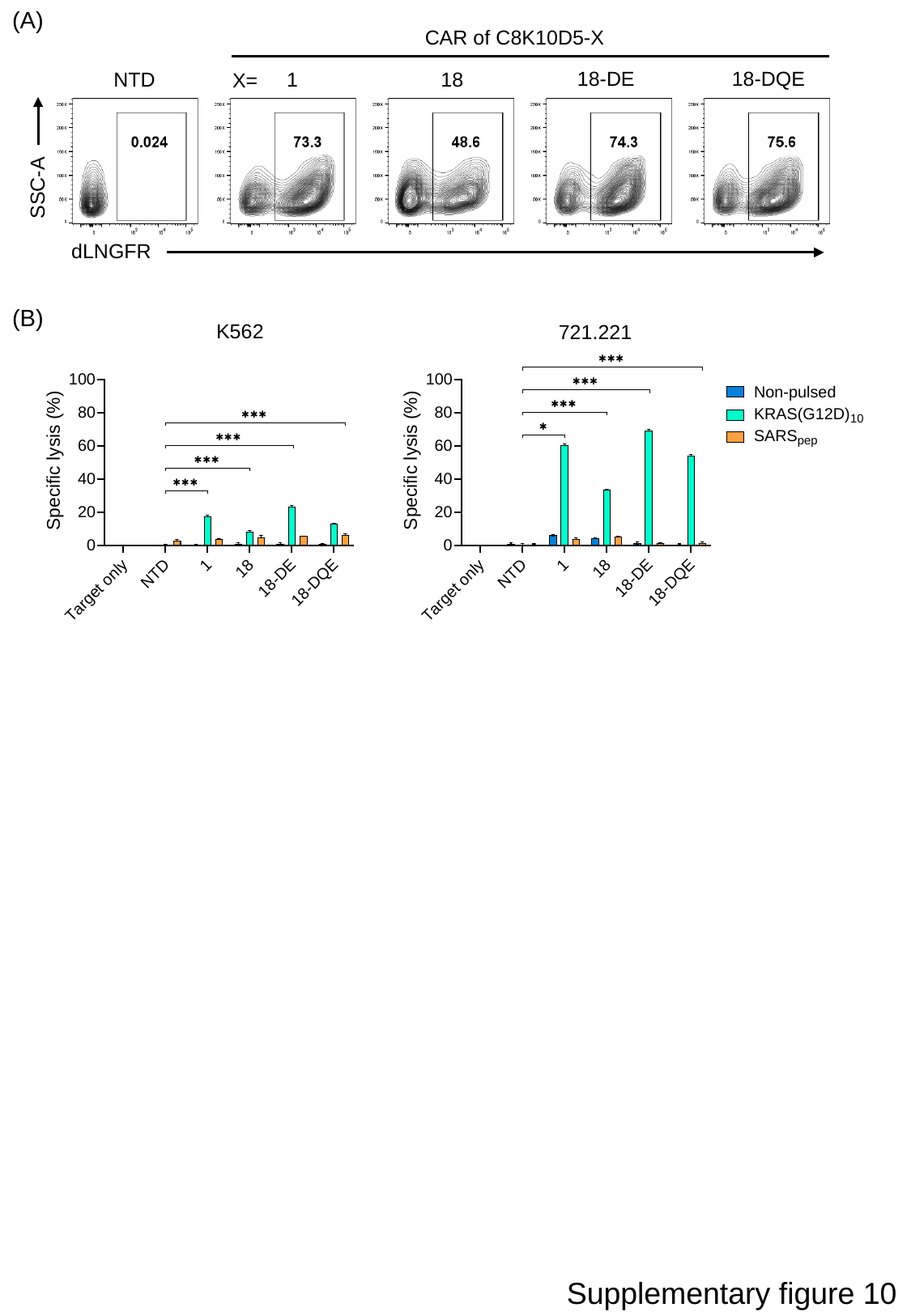

(A)
CAR of C8K10D5-X
18-DE
18-DQE
NTD
1
18
X=
SSC-A
dLNGFR
(B)
K562
721.221
Supplementary figure 10
